# Menin regulates oncogenic cell identity transcriptional networks in multiple myeloma

**DOI:** 10.64898/2026.06.17.733047

**Authors:** Emily Gruber, Subhasree Kumar, Rheana Franich, Tiffany Khong, Daniel Neville, Ching-Seng Ang, James The, Bradon Rumler, Shania Alex, Grace Dobbs, Alexander C. Lewis, George Howitt, Edwin Hawkins, P. Leif Bergsagel, Marta Chesi, Andrew Spencer, Omer Gilan, Lev M. Kats

## Abstract

Multiple myeloma (MM) is a heterogenous cancer that remains mostly incurable. A unifying feature of MM cells is a highly interconnected network of transcription factors and co-factors that coordinate the activity of super-enhancers to enforce myeloma cell identity. Targeting this network offers a promising avenue for therapeutic intervention across genetically diverse myeloma sub-types. Using integrated molecular and functional genomic approaches we identified Menin as a key driver of oncogenic gene expression in MM cells. Menin and its co-factor KMT2A bind at super-enhancers and maintain expression of essential myeloma genes, including IRF4. We demonstrate that Menin inhibitors, which have recently been approved for treatment of acute myeloid leukaemia, are highly active in myeloma cell lines and *in vivo* models. Combining Menin inhibitors with other super-enhancer targeting therapies such EP300/CREBBP inhibitors or immunomodulatory drugs (IMiDs) overcomes epigenetic plasticity in cells resistant to single agent treatment.

## Introduction

Multiple myeloma (MM) is a common plasma cell malignancy with a growing incidence in many developed nations (1). Over the past two decades, sequential introduction of novel therapies has significantly improved outcomes for MM patients, with median overall survival increasing to 5-7 years. Most recently, the introduction of T-cell redirecting therapies including CAR-T cells and bi-specific antibodies have resulted in hitherto unprecedented response rates, even in patients that had failed multiple prior lines of treatment. Nonetheless, responses are not sustained long-term in a large proportion of patients, with most ultimately experiencing relapse and progression (2,3). Hence, there is an urgent unmet need to develop novel treatment strategies, especially those that engage different mechanisms of action compared with agents currently used in the clinic.

Enhancers are distal DNA regulatory elements that bind transcription factors (TFs) and transcriptional co-activators and regulate gene expression by interacting with proximal gene promoters. Large clustered arrays of enhancers termed super-enhancers (SEs) maintain lineage-specific programs and thereby play a critical role in defining cell identity (4,5). As is the case in many cancers, MM is characterised by a highly active and interconnected SE network that sustains oncogenic gene expression. SEs within immunoglobulin heavy and light chain loci (*IGH*, *IGL*, *IGK*) are hijacked by recurrent chromosomal translocations to promote transcription of key myeloma drivers including CCND1, MAF, NSD2 and MYC. These in turn connect through reciprocal feed-forward loops to other genes such as IRF4, XBP1 and E2F that are required for MM cell survival and proliferation (6–14). A notable consequence of this aberrant transcriptional landscape is that MM cells are highly sensitive to perturbation of the epigenetic and transcriptional regulators that control it (15,16). Examples with proven clinical activity include the TFs IKZF1 and IKZF3 which bind to enhancers to upregulate IRF4 and MYC and are targeted by immunomodulatory drugs (IMiDs); and the paralogous acetyltransferases EP300 and CREBBP that broadly activate SEs in MM and are targeted by inobrodib (17–20).

Menin is a ubiquitously expressed nuclear adaptor protein that regulates gene expression through its interactions with various co-factors including KMT2A (also known as MLL1). The Menin-KMT2A complex deposits the active chromatin mark H3K4me3 and has been implicated as a positive regulator of transcription (21,22). In defined sub-types of acute leukaemia, the interaction between Menin and KMT2A is critical for the expression of self-renewal genes, most notably *MEIS1*, *MEF2C* and *PBX3*. In those settings, Menin inhibitors (iMenin) that disrupt Menin-KMT2A binding demonstrate potent anti-leukaemia activity in model systems and patients, leading to recent FDA approval for revumenib and ziftomenib in relapsed-refractory acute myeloid leukaemia (AML) (23–34). While iMenin are poised to significantly improve outcomes in AML, potential for rational deployment into other cancer types has remained underexplored. Here we demonstrate that Menin is an essential component of the myeloma cell identity circuit and provide a pathway for clinical translation of iMenin into MM by establishing response biomarkers and identifying combination strategies.

## Results

### Myeloma cells are sensitive to genetic or pharmacological disruption of Menin

To identify cancer types beyond acute leukaemias that may be sensitive to Menin inhibition we analysed the Cancer Dependency Map (DepMap) which contains pooled CRISPR screen data from >1000 cell lines (35). Stratifying cell lines by cancer lineage revealed that MM cell lines exhibit the greatest average dependency on MEN1, the gene encoding Menin (Figure 1A). MEN1 dependency in myeloma is strongly correlated with dependency on its binding partner KMT2A (Figure S1A). In a parallel analysis, MEN1 ranked among the top selective dependencies in MM (i.e. genes for which there is a greater requirement in MM compared with other cancer types), on par with the TFs IKZF1 and IKZF3 that are established anti-myeloma drug targets (Figure 1B). IRF4, well-known to be pan-essential in MM (36), was the most selective myeloma dependency. To validate the pooled CRISPR data from DepMap we used individual guides targeting MEN1 in selected MM cell lines. As expected, MEN1 was required for the proliferation and/or survival of RPMI8226 cells and to a lesser extent MM1S cells, but largely mostly dispensable in KMS11 and JJN3 cells (Figure S1B-C).

**Figure 1:**
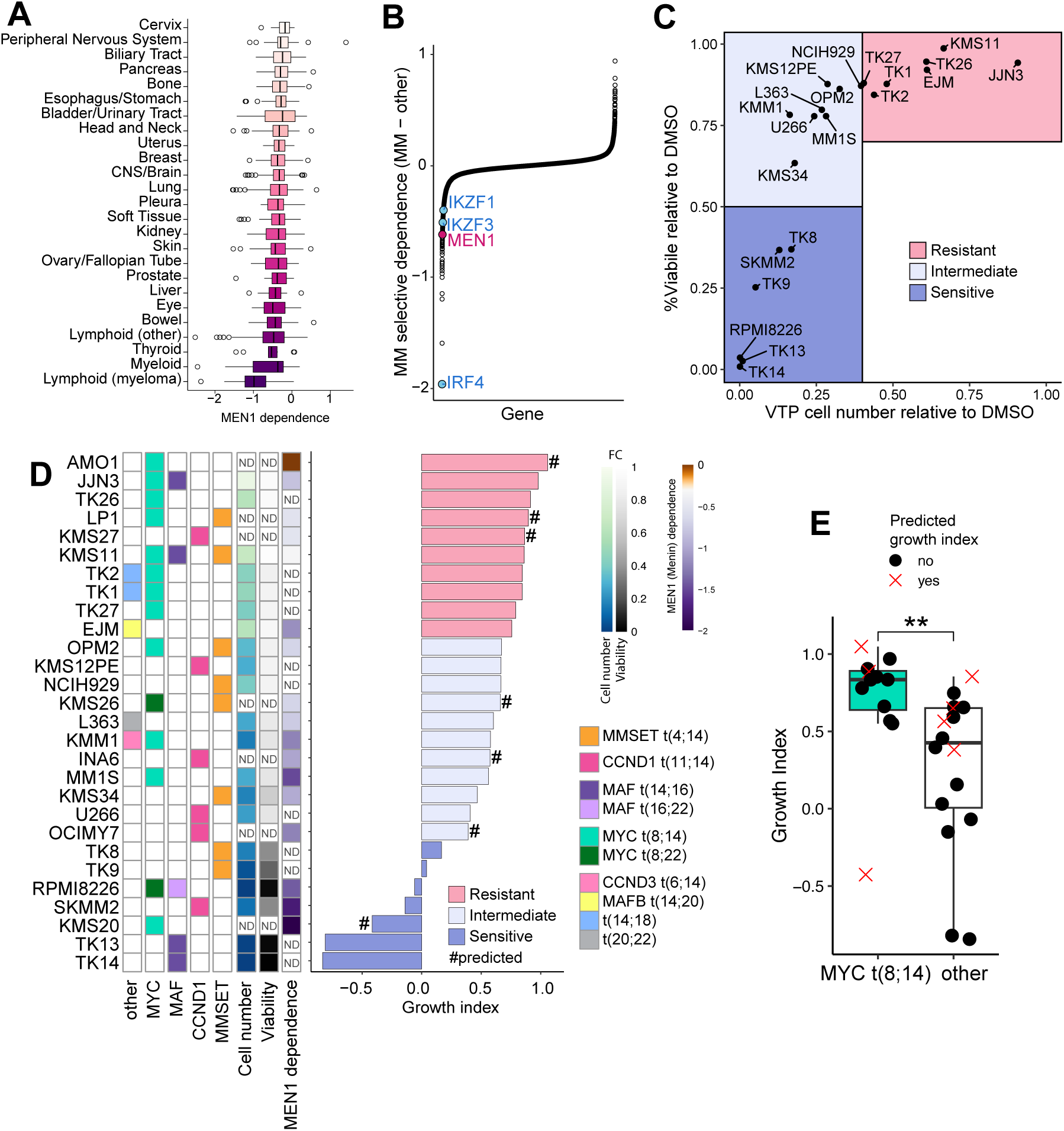
MM cell lines are sensitive to genetic and pharmacological disruption of Menin. **A.** MEN1 dependency scores across all cell lines from the Cancer Dependency Map (DepMap). **B.** Selective DepMap gene dependence in myeloma cell lines relative to all other cell lines. **C-D.** Myeloma cell lines and early-passage patient derived lines were treated with VTP-50469 (500nM) for 10 days. Cell number and viability (DAPI-) were assessed by flow-cytometry. N=3-4 biological replicates. Growth index was calculated using the GR method. Predicted growth indices were determined for cell lines with only DepMap MEN1 dependency scores. **E.** Growth index of MM cell lines stratified by the presence of t(8;14) MYC translocations.

Next, we assessed the pharmacological effects of the Menin inhibitor VTP-50469 (a close analogue of the clinical drug revumenib (27)) on a diverse panel of human myeloma cell lines, comprising 13 commercially available lines and 8 newly derived low-passage lines referred to as TK (see Methods for more information) (37). Approximately 30% (6/21) of the cell lines tested were highly sensitive to VTP-50469 treatment displaying significant viability defects (>50% cell death). An additional ∼40% (8/21) of the cell lines showed intermediate sensitivity to VTP-50469 characterised by strong anti-proliferative effects (>60% reduction in the cumulative live cell number) (Figure 1C). We determined the Menin inhibitor growth index for each cell line using the GR method (38) and observed a strong correlation between the growth effects of VTP-50469 and genetic dependence on MEN1 (Figures 1D and S1D). Notably, cell lines that are highly resistant to IMiDs such as RPMI8226 and KMM1 were responsive to VTP-50469, pointing to the potential of iMenin to overcome resistance to current standard-of-care myeloma therapies (18).

Given the variable response of MM cell lines to VTP-50469 treatment we sought to identify biomarkers that predict iMenin sensitivity. VTP-50469 responsiveness did not correlate with baseline Menin expression at the RNA or protein level, similar to what has been observed in AML cell lines (27) (Figure S1E-F). This led us to ask whether recurrent myeloma chromosomal translocations may explain differential sensitivity. To bolster the power of our analysis we devised a method to predict VTP-50469 growth index from the DepMap MEN1 dependency score. We fitted a linear model using the 10 cell lines for which both data were available, enabling us to generate a unified iMenin growth index for all 28 cell lines represented in either dataset. Correlating the unified growth index with translocation sub-types revealed that cell lines lacking the t(8;14) translocation (IGH-MYC) were enriched among iMenin sensitive and intermediate groups (Figures 1D-E and S1G). Notably however, MYC expression was not correlated with iMenin sensitivity, suggesting that the specific t(8;14) genomic event rather than MYC overexpression *per se*, is associated with reduced iMenin sensitivity (Figure S1H). Altogether, our data show that the majority of MM cell lines are dependent on MEN1 and responsive to pharmacological inhibition of Menin, supporting further investigation of iMenin as a targeted therapy in MM.

### iMenin downregulates IRF4 in responsive MM cell lines

To characterise the transcriptional consequences of Menin inhibition in MM we performed RNA sequencing (RNAseq) on RPMI8226 (iMenin sensitive), U266 (iMenin intermediate) and JJN3 (iMenin refractory) cells treated with VTP-50469 for 3 or 5 days (Figures 2A-B). The magnitude of the transcriptional response across the 3 cell lines broadly correlated with their iMenin sensitivity. VTP-50469 induced profound transcriptional changes in RPMI8226 cells after 3 days (878 differentially expressed genes (DEGs)), downregulating the essential myeloma TFs IRF4, MEF2C, PRDM1 and POU2AF1 and upregulating CD38 (the target of the therapeutic antibody daratumumab(39)) among a multitude of immune-related genes. By comparison, VTP-50469 induced fewer changes in U266 cells (86 DEGs at 3 days and 561 DEGs at 5 days) and had minimal impact on gene expression in JJN3 cells (101 DEGs after 5 days), consistent with their refractory phenotype.

**Figure 2:**
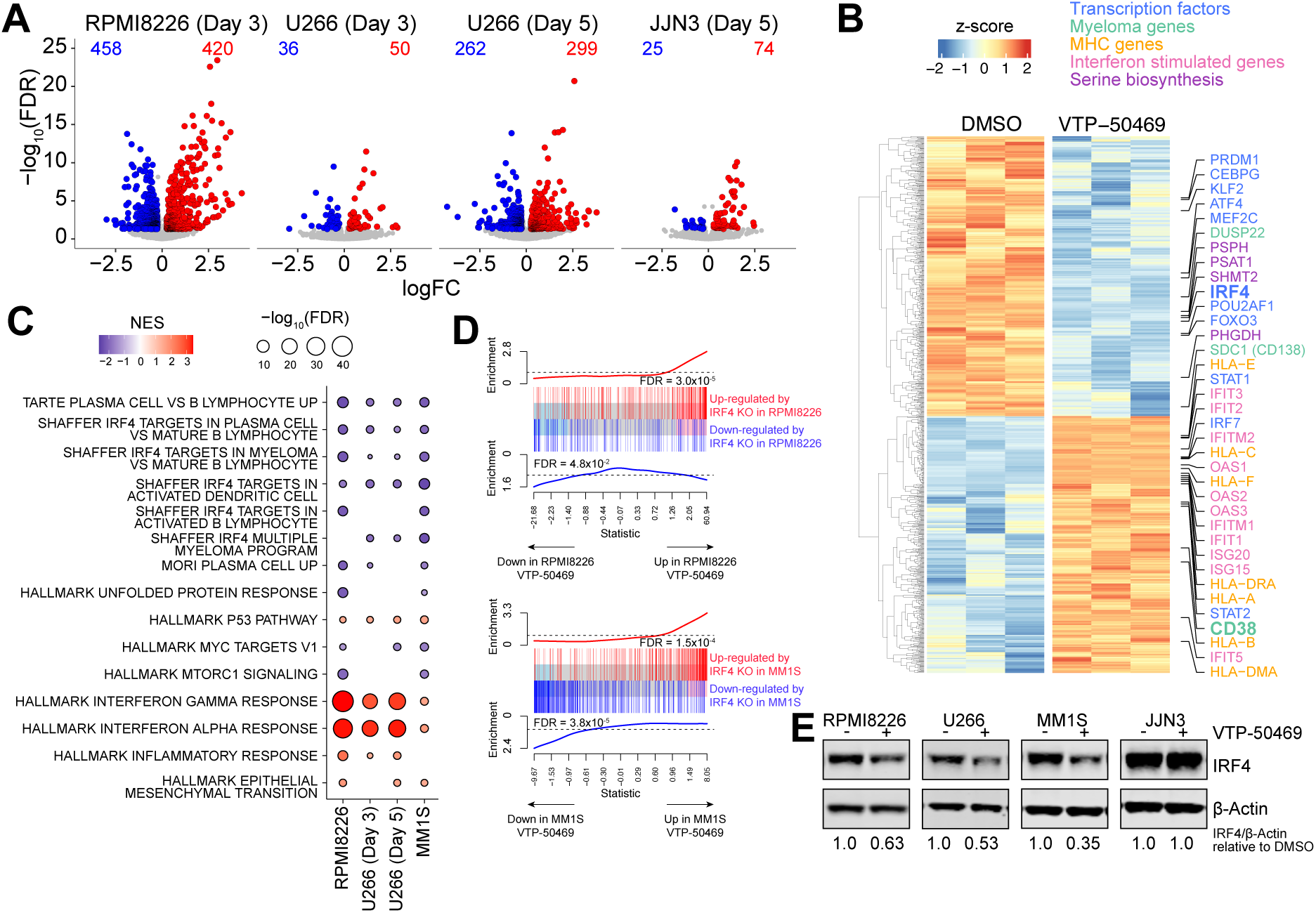
Menin inhibition downregulates core myeloma regulatory networks. **A-E.** RNAseq on myeloma cell lines treated with VTP-50469 (500nM) or DMSO for 3 (RPMI8226, U266) or 5 (U266, JJN3) days. **A.** Volcano plot highlighting down (blue) and up (red) DEGs (FDR<0.05 and |logFC|>0.25). **B.** Heatmap of DEGs in RPMI8226 cells. **C.** GSEA analysis on myeloma cell lines that are sensitive to menin inhibition, including RNAseq from MM1S cells treated with 400nM VTP-50469 for 3 days. Data plotted when p<0.1. **D.** Barcodeplots showing enrichment of gene signatures associated with IRF4 knockout (GSE148984)(40). Significance assessed using ROAST. **E.** Western blot of IRF4 following VTP-50469 treatment.

Gene set enrichment analysis in iMenin sensitive cells revealed that in RPMI8226 and U226 cells, VTP-50469 downregulated core transcriptional programs associated with myeloma proliferation and survival, including IRF4, MYC and mTORC1 (Figure 2C). These same signatures were also downregulated in another iMenin intermediate cell line, MM1S. In agreement, rotating gene set testing (ROAST) using gene sets from IRF4 knockout in RPMI8226 and MM1S cells (40) further confirmed the strong concordance with Menin inhibition, pointing to IRF4 as a major target of iMenin (Figure 2D). VTP-50469 repressed IRF4 expression at the RNA and protein levels in all iMenin responsive lines (RPMI8226, U266 and MM1S), but not iMenin refractory cells (JJN3) (Figure 2E and Table S1).

VTP-50469 treatment also led to upregulation of genes associated with interferon signalling and inflammation (Figure 2C). Of note, interferon stimulated genes (ISGs) are negatively regulated by IRF4 (Figure S2A). As expected, there was no overlap between the genes downregulated by VTP-50469 treatment in MM cells with those that have been described in AML cells (27,41), as most iMenin AML targets (e.g. MEIS1, HOXA cluster, etc) are not highly expressed in myeloma cells (Figure S2B). In contrast, there was significant overlap in the genes upregulated by VTP-50469 treatment in MM and AML cells, which predominantly encompassed ISGs and other immune genes (Figures S2B-C).

To confirm that the molecular mechanism of VTP-50469 in MM is through disruption of the Menin/KMT2A complex, we compared transcriptional changes following VTP-50469 treatment with those induced by MEN1 or KMT2A knockout. As expected, all three perturbations resulted in highly overlapping transcriptional signatures in both RPMI8226 and MM1S cells (Figures S2D-F). Overall, our data demonstrate that genetic or pharmacological perturbation of Menin disrupts oncogenic transcriptional programs in MM including those regulated by the pan-essential myeloma cell identity factor IRF4 (36).

### Menin binds the IRF4 SE

In AML cells, Menin and KMT2A co-localise at chromatin across thousands of genomic regions. Menin inhibition induces global displacement of Menin, but highly selective eviction of KMT2A, most notably at haematopoietic stem cell genes including the canonical iMenin target MEIS1 (27,41). To define the genomic occupancy of Menin and KMT2A in MM and determine how this is altered by Menin inhibition, we performed chromatin immunoprecipitation sequencing (ChIPseq). In RPMI8226, SKMM2 and MM1S cells, Menin and KMT2A occupancy were highly correlated (Figure S3A-B). Notably, there was greater enrichment of Menin at chromatin in iMenin sensitive lines RPMI8226 and SKMM2 than in the iMenin intermediate cell line MM1S, notwithstanding comparable levels of Menin protein as assessed by Western blotting (Figures S1F and S3B).

Concordant with reports in leukaemia (27,41), Menin and KMT2A localised predominantly to transcription start sites of actively transcribed genes, marked by H3K27Ac and H3K4Me3 histone modifications characteristic of active promoters (Figures 3A-B and S3B-C). VTP-50469 treatment caused global loss of Menin from chromatin, and this was accompanied by wide-spread but incomplete reduction of KMT2A. Interestingly, Menin and KMT2A also occupied many active SEs marked by broad domains of H3K27Ac but relative depletion of H3K4Me3. At these regions too, iMenin displaced Menin and reduced KMT2A.

**Figure 3:**
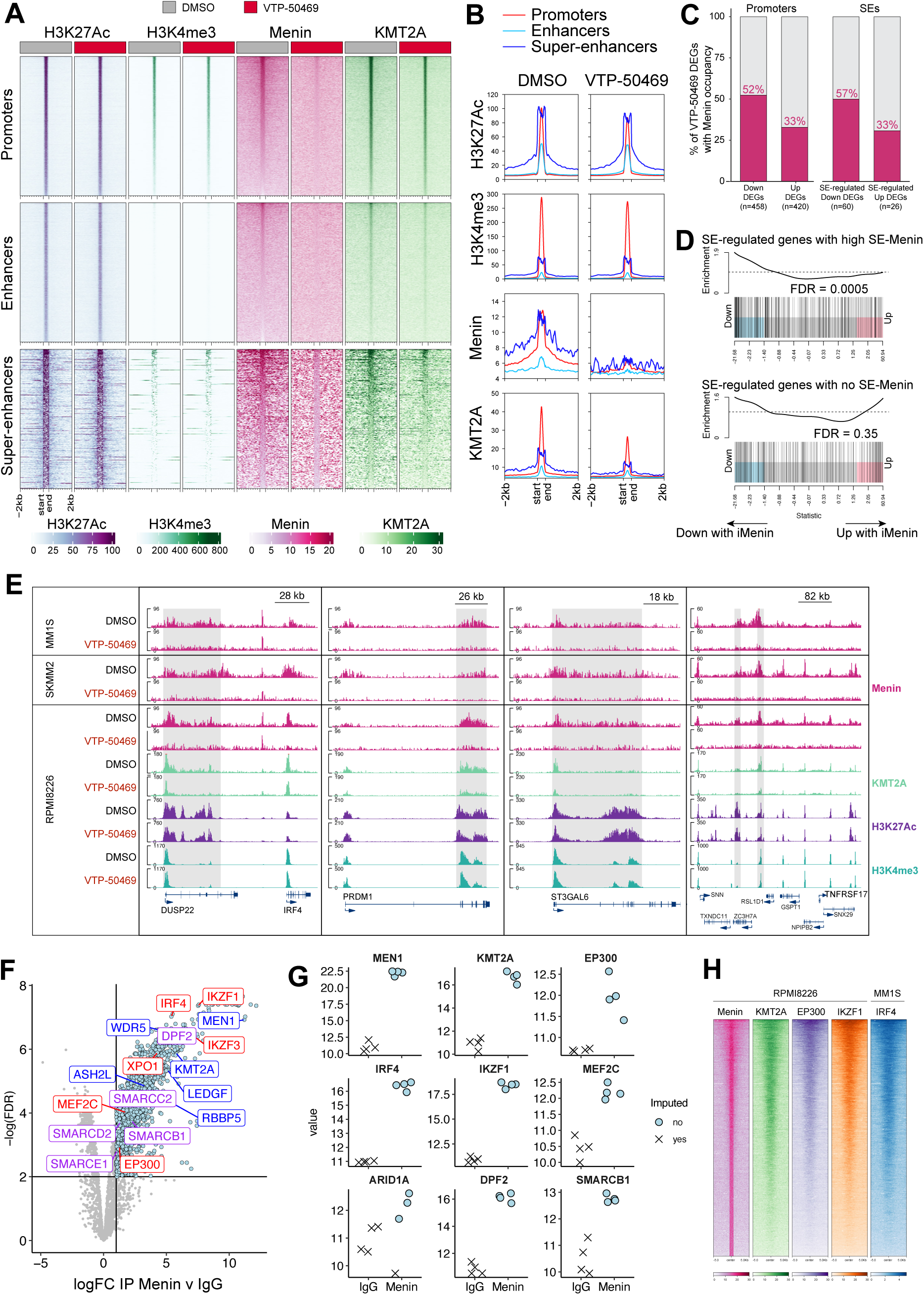
Menin/KMT2A complex is present at promoters and SEs and is evicted from chromatin by VTP-50469. **A-E.** ChIPseq in RPMI8226 cells treated with VTP-50469 (500nM) for 3 days. **A-B.** Tornado plot (A) and average signal (B) centered on peaks defined by H3K27Ac. **C.** Proportion of VTP-50469 DEGs (RPMI8226 cells) that have Menin at the promoters and SE-regulated DEGs with Menin occupancy at SEs. **D.** Barcode plot of RPMI8226 cells treated with VTP-50469 (500nM) for 3 days using genesets comprising SE-regulated genes with high (top) and no (bottom) menin occupancy at SEs. **E.** ChIPseq signal across given loci. Shaded regions denote RPMI8226 SEs. ChIPseq in SKMM2 and MM1S cells treated with VTP-50469 (500nM) for 3 days. **F-G.** RIME-MS for Menin in RPMI8226 cells. Genes highlighted in blue denote members of the Menin/KMT2A complex. Genes highlighted in red are important for transcriptional regulation in MM cells. Genes highlighted in purple are components of the cBAF SWI/SNF complex. **H.** Tornado plot centered on Menin peaks in RPMI8226 cells.

To identify direct Menin-regulated targets, we overlapped iMenin DEGs with Menin bound regions, focusing on promoters and SEs as these had the highest average Menin signal (Figure 3B). We used published H3K27Ac HiChIP data in RPMI8226 cells to assign SE-associated target genes (42). Concordant with the established role of the Menin-KMT2A complex as a transcriptional activator (21,22), iMenin downregulated DEGs were almost twice as likely to contain Menin at their promoter or associated SE as iMenin upregulated DEGs (Figure 3C). Moreover, genes associated with Menin-bound SEs were highly enriched for iMenin downregulated DEGs, whereas SEs devoid of Menin did not correlate with iMenin induced gene expression changes (Figure 3D). Menin-bound SEs (Menin-SEs) included several MM-specific regulatory regions controlling genes such as IRF4, PRDM1, SDC1 (CD138), TNFRSF17 (BCMA), IKZF1 and ST3GAL6 (43) (Figure 3E and Table S3). Additional Menin-regulated targets were enriched for metabolic processes including amino acid metabolism (Figure S3E). Altogether, these data suggest that Menin functions at SEs to sustain transcriptional networks that are critical for MM cell survival and proliferation.

To gain further insight into the interactions between Menin and other transcription and epigenetic factors we performed RIME (Rapid Immunoprecipitation Mass spectrometry of Endogenous proteins) (Figure 3F-G). In RIME, chromatin associated complexes are stabilised by formaldehyde cross-linking followed by affinity purification of the target protein and identification of its binding partners by quantitative proteomics (44). As expected MEN1 pulldown was enriched for KMT2A and other known components of the Menin-KMT2A complex including LEDGF (45). Strikingly, Menin also associated with critical myeloma transcriptional regulators including IRF4, IKZF1, IKZF3, XPO1, MEF2C, EP300 and members of the canonical BAF (cBAF) SWI/SNF chromatin remodelling complex. Consistent with these findings, examination of publicly available IRF4, IKZF1 and EP300 chromatin binding patterns in RPMI8226 and MM1S cells revealed extensive overlap with Menin/KMT2A binding, including at the Menin-SEs (Figure 3H and S3F-G). Thus, Menin is physically intertwined with the core cell identity program of MM cells.

### Genome-wide CRISPR screen identifies the EP300/NCOR1 axis as a key regulator of iMenin sensitivity

To delineate molecular pathways that regulate iMenin sensitivity in MM we performed a genome-wide CRISPR knockout screen. We reasoned that carrying out the screen in the VTP-50469 intermediate cell line MM1S would enable the identification of both sensitizer and resistance genes. MM1S cells expressing Cas9 were transduced with the Brunello sgRNA library and cultured in the presence of DMSO (control) or VTP-50469, followed by quantification of sgRNA distribution by sequencing. We then used the MAGeCK algorithm to rank enriched genes and the STRING protein-protein interaction database to collate individual hits into functional networks (46,47).

EP300 and CREBBP were among the top ranked VTP-50469 sensitiser genes (Figures 4A-B). EP300/CREBBP acetylate a range of protein targets including histones and transcription factors, and are well known dependencies in myeloma (16,48). As expected, EP300/CREBBP targeting guides were depleted with control treatment but were depleted to a greater extent with VTP-50469 treatment. Conversely, components of the NCOR complex including NCOR1 and HDAC3 that are the major antagonists of EP300/CREBBP in MM scored as strong VTP-50469 resistance hits (20). These data point to the Menin/KMT2A complex and EP300/CREBBP working in concert to maintain proliferation and survival of MM cells.

**Figure 4:**
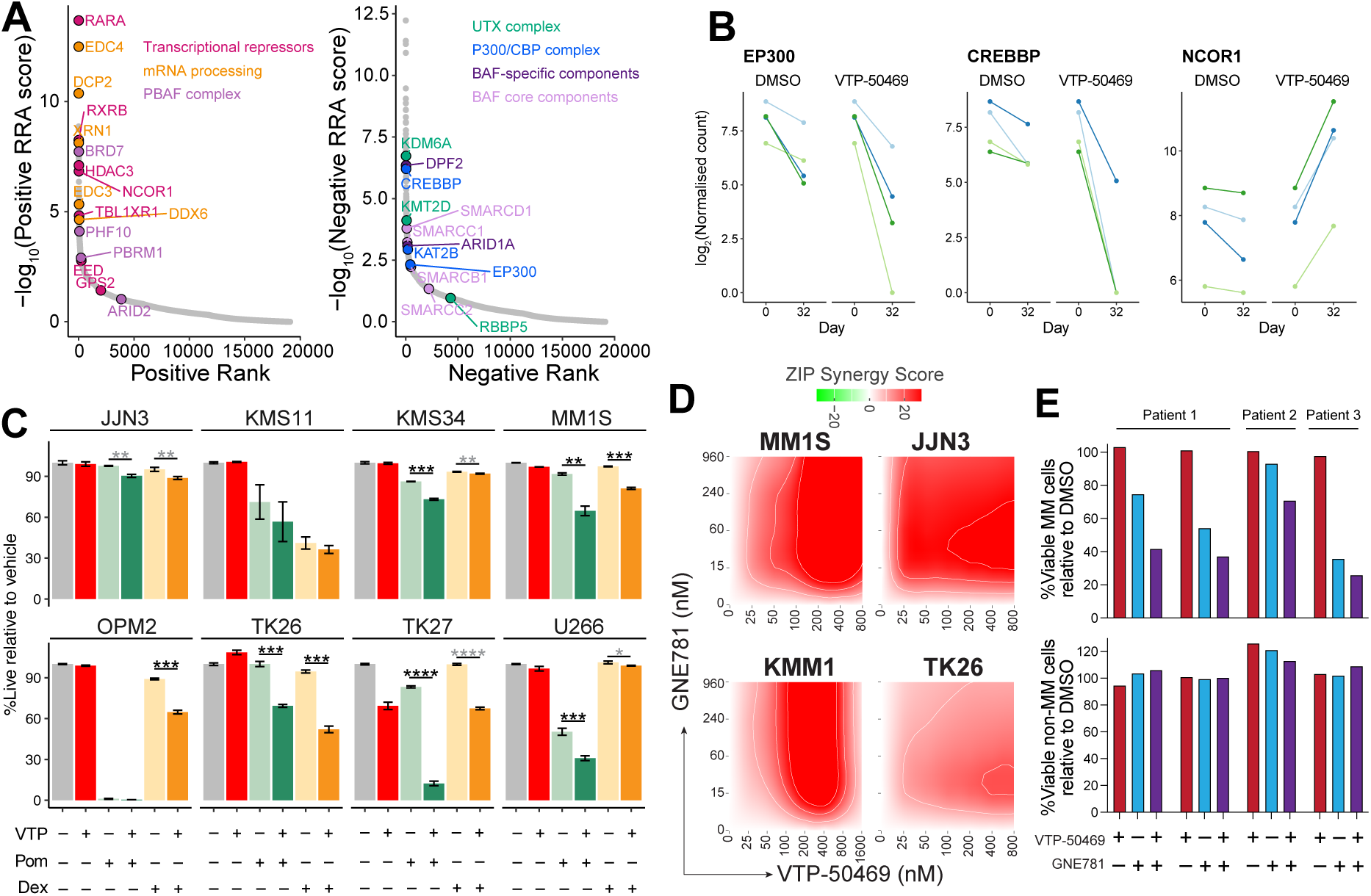
Genome-wide CRISPR screen identifies the EP300/NCOR1 axis as a key regulator of iMenin sensitivity. **A-B.** CRISPR screen in MM1S cells cultured for 32 days in increasing concentrations of VTP-50469 (200-400nM). **A.** MAGeCK RRA scores for genes that confer resistance (left) and sensitization (right) to VTP-50469. **B.** sgRNA plots. **C.** Myeloma cell lines treated for 7 days with VTP-50469 (VTP; 500nM) in combination with standard of care therapeutics pomalidomide (Pom; 400nM) and dexamethasone (Dex; 20nM). Viability assessed with DAPI staining by flow-cytometry. N=3 biological replicates. Significance assessed by t-test: ****p<0.0001, ***p<0.001, **p<0.01, *p<0.05. Stars colored black have a bliss independence synergy score > 10 and grey when the bliss independence synergy score ≤ 10. **D.** Synergy score calculated using synergyfinder on cells treated with VTP-50469 and the EP300/CREBBP bromodomain inhibitor GNE781 over 4 days. DAPI+ cells were quantified by flow-cytometry. N=3 biological replicates. **E.** Primary patient samples were treated *ex vivo* with VTP-50469 (500nM) and/or GNE781 (960nM) for 72h and flow-cytometry used to quantify viable MM (CD45-/CD38+/Apo2.7-) or non-MM (CD45+/CD38-/Apo2.7-) cells. Samples from patient 1 were collected 2 months apart.

Multiple members of the SWI/SNF family of ATP-dependent chromatin remodelling complexes also scored in our screen (Figures 4A and S4A). SWI/SNF complexes are highly modular, with shared core units assembled with different specialised subunits to form cBAF, PBAF and ncBAF complexes with distinct localisation patterns and function (49,50). cBAF complexes primarily occupy and maintain active distal enhancers, whilst PBAF complexes typically localize to active promoters. While BAF core components (SMARCD1, SMARCC1/2, SMARCB1) and cBAF-specific units (ARID1A, DPF2) were identified as sensitiser hits, PBAF components (BRD7, PHF10, PBRM1), scored as resistance hits. Notably, recent work has implicated cBAF as a key driver of IRF4-dependent transcriptional networks (51).

The KDM6A (aka UTX) complex also emerged as an iMenin sensitiser. KDM6A is an H3K27-specific demethylase that binds KMT2D (or KMT2C) and is involved in enhancer rewiring (52). KDM6A is mutated in ∼5% of MM patients and associated with poor survival (53). Deletion of KDM6A or KMT2D was recently reported to mediate resistance to Menin inhibition in AML, pointing to divergent interplay of Menin-KMT2A and KDM6A-KMT2D complexes between AML and MM contexts(54). Additionally, disruption of mRNA stability control was implicated in driving iMenin resistance with numerous genes involved in mRNA decapping (DCP2, EDC4, XRN1, DDX6) scoring as VTP-50469 resistance hits (55). We further independently validated selected positively and negatively enriched genes using proliferative competition assays (Figure S4B). Collectively, these findings further highlight Menin as a driver of the oncogenic regulatory network in MM and uncover pathways that influence the potency of Menin inhibition.

### Combinatorial activity of iMenin with established and emerging anti-myeloma drugs

To enhance the translational relevance of our findings, we next evaluated the effects of combining iMenin with other drugs that are in clinical use or advanced development in myeloma. First, we tested VTP-50469 with the corticosteroid dexamethasone and the iMiD pomalidomide, both of which are extensively used in front-line myeloma regimens (2). Of note, iMenin and IMiDs have demonstrated combinatorial activity against pre-clinical AML models (56,57). We focussed our analysis on iMenin intermediate and refractory MM cell lines (Figure 4C). In most of these lines, 7 days of single agent dexamethasone or pomalidomide treatment had a minimal cytotoxic effect and the addition of VTP-50469 increased the killing by either or both drugs. The effects were most pronounced in MM1S, TK26, TK27 and U266 but were mild in some other lines such as JJN3.

Given our CRISPR data and the promising activity of the EP300/CREBBP bromodomain inhibitor inobrodib in ongoing trials (19,58), we next tested the combination of iMenin with EP300/CREBBP inhibitors (iEP300). As predicted by our functional genomic screening, VTP-50469 and iEP300 were significantly more potent when used together (Figures 4D and S4C-E). Excitingly, the combination was synergistic as assessed by the Zero Interaction Potency (ZIP) score (59) and extended across all cell lines analysed (including JJN3 and TK26 cells which are refractory to VTP-50469) as well as across both catalytic and bromodomain iEP300 chemotypes. The combination of iMenin and iEP300 also increased myeloma cell death by at least 25% percent in 4/4 primary patient samples (collected from 3 different patients) cultured *ex vivo*, with no impact on viability of autologous lymphocytes (Figure 4E).

### iMenin and iEP300 synergize by supressing SE networks

To understand the molecular mechanisms that underpin the observed synergy between iMenin and iEP300 we performed RNAseq in MM1S cells treated with VTP-50469, the bromodomain iEP300 GNE781, or the combination. DEGs triggered by VTP-50469 or GNE781 were highly correlated, although the extent of the transcriptional perturbation was greater in GNE781 treated cells (Figures 5A and S5A). Importantly, gene expression changes in cells treated with either single agent were amplified in the combination treatment, including marked repression of key MM TFs (*IRF4*, *MYC*, *XBP1* and *IKZF1*) and downregulation of cell-cycle and proliferation associated signatures (Figures 5A-C and S5A-B).

**Figure 5:**
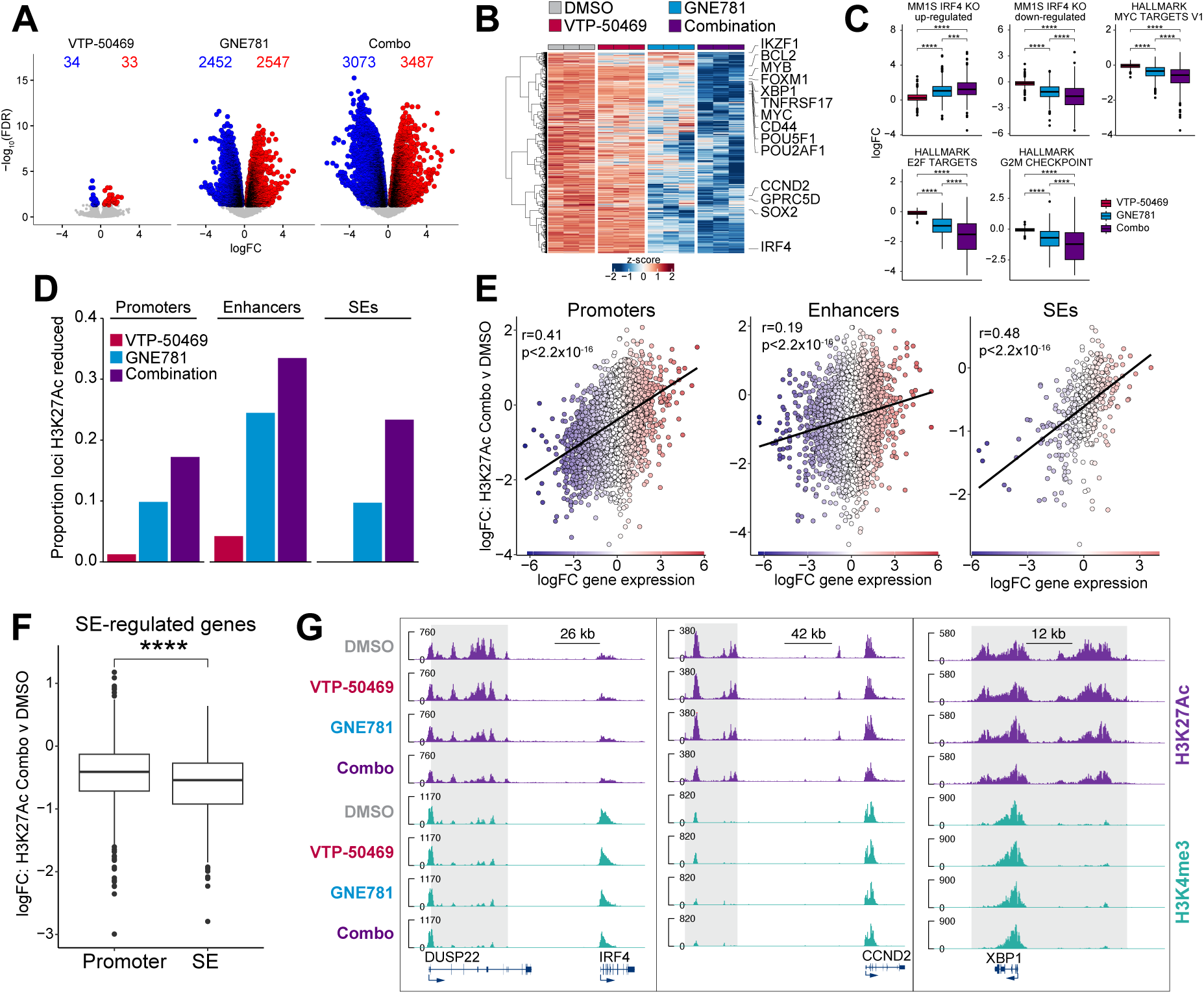
iMenin and iEP300 synergize by suppressing SE networks. **A-G.** RNAseq and ChIPseq of MM1S cells treated with VTP-50469 (400nM), GNE781 (60nM) or the combination for 3 days. **A.** Volcano plot highlighting DEGs (FDR < 0.05 and logFC < -0.25). **B.** Heatmap of combination down DEGs (FDR < 0.05 and logFC < -0.25). **C.** LogFC of the genes in given genesets. **D.** Proportion of loci that lose H3K27Ac (logFC < -1). **E.** Correlation between H3K27Ac loss and gene expression changes following combination treatment. **F.** Change in H3K27Ac at the promoters and SEs of SE-regulated genes. **G.** ChIPseq signal tracks across given loci. Shaded regions denote SEs in MM1S DMSO cells. Significance assessed with Student’s t-test. ****p<0.0001.

Next, we profiled the impact of drug treatment on the distribution of KMT2A, H3K4Me3 and H3K27Ac at chromatin (Figures 5D and S5C). As expected, VTP-50469 reduced KMT2A recruitment, whereas GNE781 preferentially decreased H3K27Ac at enhancer regions, consistent with other bromodomain and catalytic iEP300 (19,20). Relative to the single agents, the combination resulted in a pronounced reduction of KMT2A from promoters and was accompanied by a modest decrease in H3K4Me3, although this did not correlate with transcriptional changes on a genome-wide level (Figures S5C-D). The combination also resulted in greater loss of H3K27Ac across all genomic features, with a larger proportion of enhancers and SEs displaying reduced H3K27Ac in comparison to promoters (Figure 5D). Loss of H3K27Ac at SEs and promoters correlated most strongly with transcriptional repression, whilst the correlation was weak at typical enhancers (Figure 5E). Notably, at SE-regulated genes, such as *IRF4*, loss of H3K27Ac was consistently greater at the SE compared with the promoter (Figures 5F-G). The combination of iMenin and iEP300 also substantially reduced IRF4 and MYC levels in an iMenin refractory cell line (JJN3) (Figure S5E). Collectively, these data demonstrate that combined Menin and EP300/CREBBP inhibition effectively disrupts SE-associated gene networks in MM.

### The clinically approved iMenin revumenib demonstrates anti-myeloma efficacy *in vivo*

Revumenib was the first iMenin to be approved for the treatment of relapsed/refractory AML (29). To assess its anti-myeloma efficacy *in vivo,* we evaluated revumenib in an orthotopic RPMI8226 xenograft model. RPMI8226 cells expressing a luciferase reporter were engrafted into NOD-*scid* IL2Rgamma^null^ (NSG) mice by intrafemoral injection and revumenib was delivered by oral gavage at a dose of 50mg/kg b.i.d which is comparable with regimens used in pre-clinical AML models. Serial bioluminescence imaging demonstrated that 3 weeks of revumenib treatment (the maximum allowable time under our approved animal ethics protocol) induced tumour regression in all (4/4) treated mice (Figures 6A and S6A). Revumenib treatment also significantly extended overall survival compared with vehicle treated controls, although revumenib-treated animals progressed after treatment withdrawal (Figure 6B).

**Figure 6:**
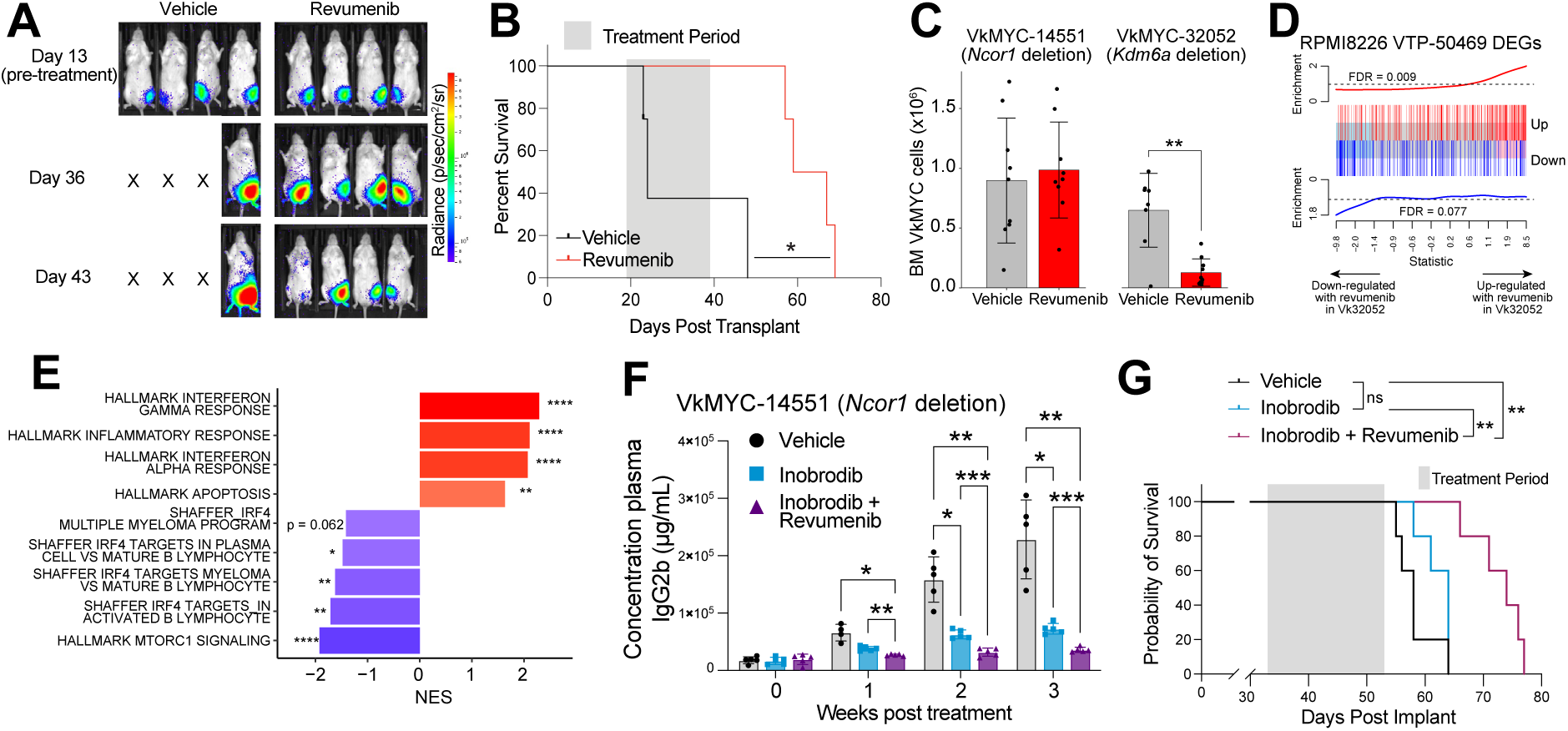
Revumenib demonstrates anti-myeloma efficacy *in vivo* as a single agent and in combination with inobrodib. **A-B.** RPMI8226 cells expressing luciferase were transplanted by intrafemoral injection into NSG mice and treated with 50mg/kg Revumenib by oral gavage b.i.d. for 3 weeks. **A.** Luciferase imaging. **B.** Kaplan-Meier analysis. Log-rank test was used to assess significance. **C-E.** VkMYC cells with a monoallelic *Ncor1* deletion (#14551) or biallelic *Kdm6a* deletion (#32052) were transplanted into C57BL/6 mice by tail vein injection and mice treated with 50mg/kg Revumenib by oral gavage b.i.d. for 3 weeks. **C.** Tumour burden per femur (VkMYC-14551) and per femur/tibia (VkMYC-32052) was quantified by flow-cytometry. Significance assessed using Wilcoxon test. **D-E.** VkMYC-32052 cells were isolated by FACS from the bone marrow of mice after 3 weeks of revumenib treatment and subjected to RNAseq. **F.** Barcodeplot showing RPMI8226 VTP-50469 DEGs are enriched in revumenib treated VkMYC-32052 cells. **G.** GSEA analysis. **F-G.** VkMYC #14551 transplanted mice were treated by oral gavage with 10 mg/kg Inobrodib q.d. with or without revumenib (50mg/kg b.i.d) for 3 weeks. **F.** ELISA for plasma IgG. Significance assessed by Student’s t-test. **G.** Kaplan-Meier analysis. Log-rank test was used to assess significance. ****p<0.0001, ***p<0.001, **p<0.01, *p<0.05

We next turned to the Vk*MYC syngeneic myeloma model which demonstrates high biological fidelity to the human disease and drug responses that are predictive of clinical efficacy (60–62). We performed 2 separate therapy experiments using 2 different transplantable Vk*MYC tumours with distinct co-operating mutations – Vk14551 which carries monoallelic deletions of *Ncor1, Cdkn2*a and *Cdkn2b*; and Vk32052 which has biallelic loss of *Kdm6a.* NCOR1 was identified as an iMenin resistance gene in our MM1S CRISPR screen, whereas KDM6A was identified as a sensitizer gene (Figure 4A). Consistent with the CRISPR screen data, revumenib treatment demonstrated efficacy in the Vk32052 model, but not in the Vk14551 model. In mice bearing Vk32052 tumours, revumenib reduced myeloma burden by ∼50% (Figures 6C and S6B-C). This was accompanied by changes in the immune compartment, with a significant decrease in macrophages, Ly6c-low monocytes and neutrophils, and a concomitant increase in CD8 T-cells in the bone marrow of revumenib treated mice relative to controls (Figures S6D-F). RNAseq on Vk32052 cells isolated from the bone marrow revealed transcriptional changes concordant with those observed in VTP-50469 treated human MM cells *in vitro*, including the downregulation of TFs *Mef2c* and *Xbp1* and gene signatures related to myeloma proliferation and survival, and an up-regulation of interferon signatures (Figures 6D-E and S6G). Although *Irf4* was not significantly downregulated, *Irf4* target genes were altered upon revumenib treatment (Figures 6E and S6H). By comparison, revumenib had no significant impact on either tumour burden or the immune milieu in mice engrafted with Vk14551 tumours (Figures S6B-F). Altogether, these studies confirm that revumenib demonstrates *in vivo* single agent anti-myeloma efficacy, at least in some contexts. They also suggest that NCOR1 and KDM6A may serve as biomarkers of iMenin resistance and sensitivity, respectively, although future testing with additional models is required to assess the generalisability of our observations.

As Vk14551 tumours were resistant to revumenib, we then asked whether the combination of revumenib and inobrodib would provide a therapeutic benefit. Single agent inobrodib treatment slowed tumour progression of Vk14551 tumours, but this resulted in only a minimal extension in overall survival that did not reach statistical significance (Figure 6F-G). By comparison the combination of revumenib and inobrodib was well tolerated and arrested tumour while the mice remained on treatment, and significantly extended survival (Figure 6F-G and S6I-J).

## Discussion

Drug resistance remains a major challenge in MM and novel strategies are required for patients that have exhausted available treatment options (2,3). By analysing gene dependency data in DepMap, we identified Menin as a promising drug target for MM. Menin is an epigenetic adaptor protein that interacts with various binding partners and regulates gene expression in a context dependent manner (21,22). Although originally discovered as a tumour suppressor in the familial cancer syndrome multiple endocrine neoplasia-type 1, subsequent work demonstrated that in AML it functions as an oncogenic co-factor (23–25,33,34,63). Efforts to develop iMenin recently paid dividend, with large clinical trials demonstrating excellent potency and tolerability in AML, leading to FDA approval (29–31). Notably however, DepMap suggested that MM cells may be even more sensitive to perturbation of the Menin-KMT2A complex than AML cells, prompting us to evaluate Menin targeting in myeloma.

Our mechanistic data implicate the Menin-KMT2A complex as an integral component of the MM core oncogenic gene regulatory network. While most studies in leukaemia have focussed on the function of Menin-KMT2A at proximal promoters, we found that in MM the complex is enriched at both promoters as well as SEs. Menin physically interacts with SE-associated lineage defining TFs including IRF4, MEF2C and IKZF1/3; and functionally co-operates with epigenetic factors that are known to maintain enhancer activation, specifically EP300/CREBBP, cBAF and KDM6A-KMT2D (17,18,20,50,52). MM has been referred to as an “enhanceropathy”, with expression of key myeloma drivers commonly driven by highly active plasma cell or myeloma-specific Ses (13,14). Our findings suggest Menin contributes to this aberrant epigenetic landscape, and that iMenin collapses key nodes of the transcriptional circuit required for MM survival and proliferation.

Assessing iMenin sensitivity across a panel of MM cell lines, we observed a range of responses with an overall response rate of approximately 60%. MM is a highly heterogenous cancer, so the variable sensitivity was expected and predicted by DepMap. The range of responses to iMenin is similar to that of pomalidomide and lenalidomide, drugs with proven clinical efficacy in MM (17,18). Absence of the t(8;14) chromosomal translocation was the best genomic predictor of iMenin sensitivity, although our cell line panel had limited power to identify genomic biomarkers. The sensitivity of different MM cells to iMenin was broadly correlated with the impact of iMenin on gene expression. In highly sensitive cell lines such as RPMI8226, transcriptional changes induced by iMenin occurred more rapidly and were of a greater magnitude compared to cell lines with intermediate iMenin sensitivity where the changes were more modest and took longer. These observations are in line with what has recently been reported with iMenin in leukaemia (64,65) and with IMiDs in myeloma (17,18), and likely reflect context-specific wiring of gene regulatory networks and their capacity to withstand loss of individual components.

An attractive strategy to combat the plasticity of the MM oncogenic network is to combine multiple agents that target its different nodes. Our findings demonstrate that iMenin can potentiate the activity of other agents that target MM SEs including IMiDs and iEP300. Of the combinations tested, dual Menin and EP300/CREBBP inhibition was the most synergistic and demonstrated potent activity against primary patient samples and in the highly predictive syngeneic Vk*MYC model (60). Importantly, unlike other co-activator targeting combinations such as GNE781 and the BRD4 inhibitor JQ1 that are highly toxic, the combination of revumenib and inobrodib was well tolerated in mice. Whether it is possible to boost efficacy with triplet combination of iMenin, iEP300 and IMiDs, or through addition of emerging agents such as SMARCA2/4 inhibitors (66), remains to be seen. Moreover, the induction of ISGs including CD38 suggests potential for combining iMenin with daratumumab that will need to be assessed in future studies.

Altogether, we believe that our study provides compelling evidence for prioritising iMenin for clinical testing in MM. Development of multiple clinical-grade iMenin over the past 5 year and the potential to combine them with drugs already approved for myeloma opens the path for translation of our research into improving treatment for myeloma patients.

## Data availability

Processed and raw RNAseq and ChIPseq data have been deposited to GEO and will be made available upon publication. Raw counts and outputs from MAGeCK for the CRISPR screen can be found in Supplementary Table S5. Differential gene expression tables are available in Supplementary Tables S1-2,7-8.

## Author disclosures

The authors have no conflicts to disclose.

## Author contributions

L.M.K., E.G. and O.G. designed the study, wrote the manuscript and supervised the study; E.G., S.K., R.F., T.K., D.N., C.A., J.T., B.R., S.A., G.D. and A.C.L. performed the experiments and analyzed the data; T.K. G.H., E.H., P.L.B., M.C. and A.S. provided technical expertise and essential reagents.

## Acknowledgements

We thank the members of the molecular genomics, animal and flow cytometry core facilities at the Peter MacCallum Cancer Centre and the Peter MacCallum Cancer Centre Models of Cancer Translational Research Centre for technical support; and members of the Barrie Dalgleish Centre for Multiple Myeloma and Related Blood Cancers for advice and discussion. This work was supported by grants from the National Health and Medical Research Council of Australia, the Barrie Dalgleish Foundation and the Peter MacCallum Cancer Centre Foundation. S.K. was supported by the Australian Postgraduate Award. L.M.K is supported by the Blood Cancer United (formerly Leukemia and Lymphoma Society) Scholar Award.

## Methods

### Cell culture

MM cell lines were verified by short tandem repeat profiling and routinely tested for mycoplasma contamination. MM cells were cultured at 37°C and 5% CO_2_ in culture medium outlined in Table 1. HEK293T cells were cultured in DMEM (Gibco) supplemented with 10% FBS and 1% penicillin/streptomycin.

**Table 1.** Cell culture conditions.

| CELL LINE | MEDIA |
| --- | --- |
| SKMM2, KMS-12-PE | RPMI-1640 (Gibco)+ 20 % FBS + 1% penicillin/streptomycin |
| EJM | IMDM + 15 % FBS + 1 % penicillin/streptomycin |
| L363 | RPMI-1640 (Gibco) + 15 % FBS + 1 % penicillin/streptomycin |
| NCI-H929/H929 | RPMI-1640 (Gibco) + 10 % FBS + 50 uM 2-ME + 1 % penicillin/streptomycin |
| KMS-11, KMS-34, RPMI, MM1.S, OPM2, MM1.R, U266 | RPMI-1640 (500 mL) + 10 % FBS (50 mL) + 1 % penicillin/streptomycin |
| JJN3 | RPMI-1640 (500 mL) + 10 % FBS (50 mL) + 5 mL GlutaMax + 5 mL Sodium Pyruvate + P/S (2.5 mL) |
| KMM1 | RPMI-1640 (500 mL) + 10 % FBS (50 mL) + 5 mL GlutaMax + P/S (2.5 mL) |
| EPPDs/TK cell lines | RPMI-1640 (500 mL) + 10 % FBS (50 mL) + 1 % penicillin/streptomycin + 10 % conditioned media (HS5 stromal media) for one week, then cultured without conditioned media |
| HEK293T | DMEM (500 mL) + 10 % FBS (50 mL) + 1 % penicillin/streptomycin |

### Animal experimental models

All animal experiments were performed at the Peter MacCallum Cancer Centre in an approved premises nominated on the Bureau of Animal Welfare Scientific Licence SPPL20183 (Agriculture Victoria, Australia) and were approved by the Peter MacCallum Cancer Centre Animal Experimentation Ethics Committee. Female C57BL/6 and immune deficient NOD.Cg-Prkdcscid Il2rgtm1Wjl/SzJ (referred to as NSG) mice were bred within the Peter MacCallum Cancer Centre animal facility or purchased from the Walter and Eliza Hall Institute of Medical Research (Melbourne, Australia). All experimental mice were housed at the Peter MacCallum Cancer Centre animal facility under specific pathogen-free conditions. Animals were group-housed in individually ventilated micro-isolator cages (6 mice per cage) on a 13hr light/11-hr dark cycle. Mice had continuous access to sterilized water and Barastoc irradiated mouse cubes (Ridley AgriProducts).

### Primary MM samples

Primary multiple myeloma samples were obtained from patients with relapsed/refractory multiple myeloma following written informed consent and approval from the Alfred Hospital Research and Ethics Committee (Project 29/05). Bone marrow mononuclear cells (BMMNCs) were isolated using Ficoll-Paque Plus (Amersham Biosciences), washed in PBS, and subjected to red blood cell lysis buffer using NH4Cl solution (8.29 g/L ammonium chloride, 0.037 g/L ethylenediaminetetraacetic acid [EDTA], and 1 g/L potassium bicarbonate). Following additional PBS washes, BMMNCs were quantified using a TC20 automated cell counter (Bio-Rad) and cultured in RPMI-1640 medium supplemented with 10% heat-inactivated foetal bovine serum (FBS) and 2 mmol/L L-glutamine.

### Cloning

sgRNAs were cloned into lentiviral vectors FgH1tUTG-GFP (Addgene plasmid #70183). sgRNA sequences are provided in Table 2. sgRNA oligos were annealed by step-down PCR in T4 ligase buffer (NEB) prior to ligation into Esp3I cut FgH1tUTG or lentiguide-Crimson using T4 ligase according to manufacturer’s protocols. Ligated products were transformed in DH5a competent bacteria using standard heatshock protocols. Single bacteria colonies were picked to validate by sanger sequencing for validation prior to transfection into 293T cells for lentiviral transduction. MSCV-GFP-IRES-Luc2 was a gift from the Johnstone Laboratory.

**Table 2.**
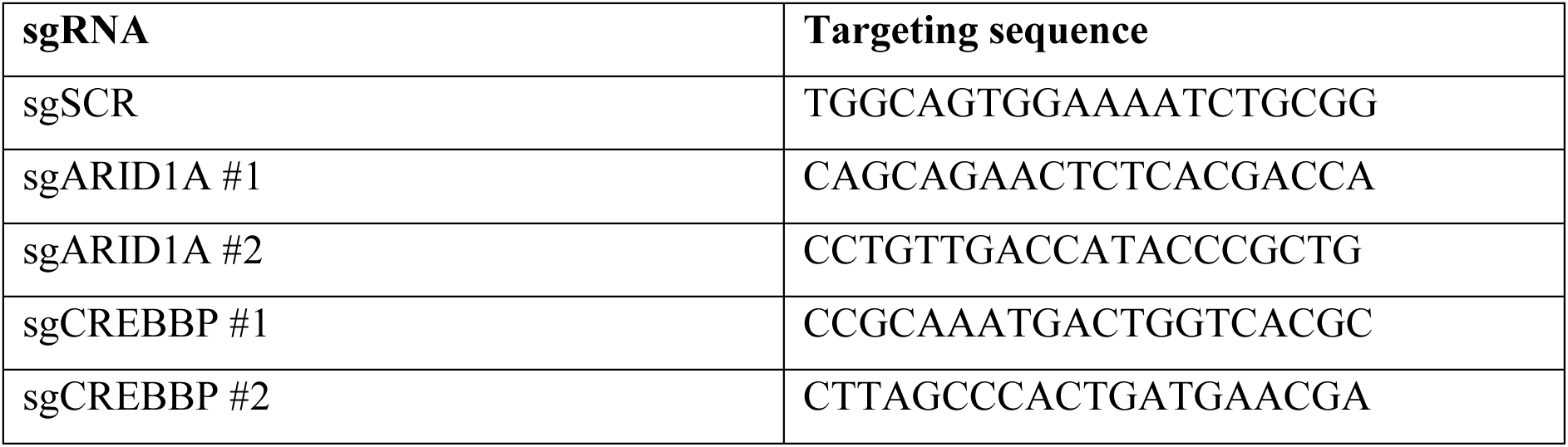
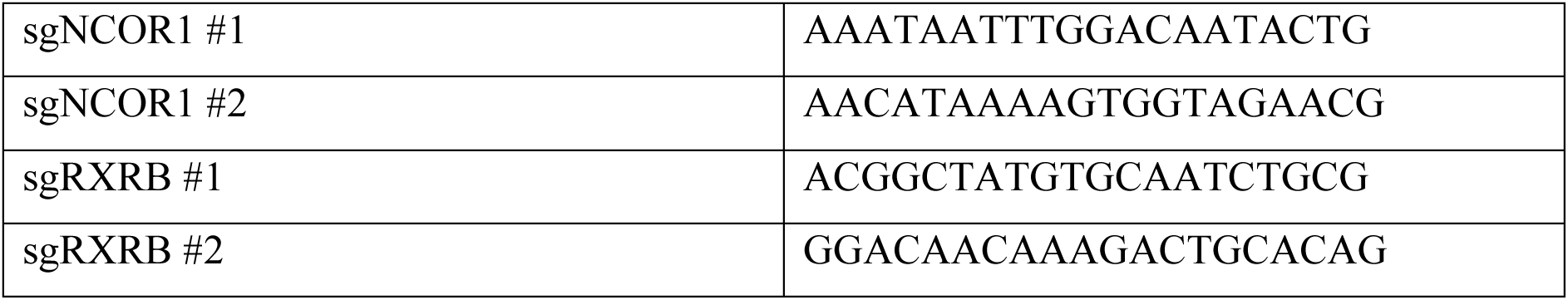
sgRNA sequences.

### Viral transduction

Non-replicating lentiviruses were generated by transient co-transfection of the transfer plasmids into HEK293T cells together with the packaging plasmids pMDL (Addgene plasmid #12251), pRSV-REV (Addgene plasmid #12253), and pVSVG (Addgene plasmid #12259) with PEI (1µg/mL). Viral supernatant was collected 48h post transfection and passed through a 70µm filter. Transduction was performed by centrifuging target cells with viral supernatant at 1000 xg in the presence of 4 µg/ml sequabrene (Sigma-Aldrich).

To generate MSCV-GFP-IRES-Luc2 RPMI8226 cells, 15x10^6^ HEK293T cells were plated in a T175 flask 24h prior to co-transfection with MSCV-GFP-IRES-Luc2 and the ecotropic packaging vector pCL–Ampho. Viral supernatant was collected at 48h post transfection, filtered, concentrated (Amicon Ultra 15mL 100K #UFC910008) and spun onto Retronectin (T100A)-coated 6 well plates at 2000xg for 1h. 5x10^5^ RPMI8226 cells were spun onto a virus coated well at 1400rpm for 5mins and cultured at 37°C for 24h.

### RPMI8226 xenograft

Approximately 1x10^6^ RPMI8226 cells transduced with the MSCV-GFP-IRES-Luc2 plasmid and transplanted into NSG mice by intrafemoral injection. Following an initial 18-day engraftment period mice were randomized to receive Revumenib (50mg/kg) or vehicle. Drug was administered twice daily by oral gavage for 5-days followed by once daily for 2-days over a 3-week period. Animals were closely monitored for clinical signs of disease development (weight loss, lethargy, hunched posture) and tumor burden measured weekly by luciferase imaging. Mice were administered 50mg/kg D-luciferin (Revivify, #122799A) by IP injection and anesthetized by isoflurane 5min after injection. Bioluminescence was imaged using the IVIS100 imaging system (Perkin Elmer) over a 20sec and 60sec period. Living Image (v 4.8.0) was utilized to analyze images and quantify bioluminescence. Animals were euthanized at ethical endpoint based on clinical symptoms and overall survival rates were assessed using Kaplan-Meier analysis.

### VkMYC *in vivo* experiments

VkMYC-32052 cells were kind gifts from Prof Marta Chesi. VkMYC-14551 cells that had been modified by transduction to express GFP were kindly provided by Prof Edwin Hawkins. 0.25x10^6^ VkMYC-14551 and VkMYC-32052 cells were transplanted into C57Bl/6 mice by tail vein injection. Following an initial engraftment period (3d VkMYC-32052 and 33d VkMYC-14551) mice were randomized to receive Revumenib (50mg/kg) or vehicle. Drug was administered by oral gavage twice daily for 5-days followed by once daily for 2-days over a 3-week period. For the combination experiment, mice received vehicle, Revumenib (50mg/kg), Inobrodib (10mg/kg) or the combination of Revumenib (50mg/kg) and Inobrodib (10mg/kg). Revumenib was administered as described above and Inobrodib was administered q.d. by oral gavage. Animals were closely monitored for clinical signs of disease development (weight loss, lethargy, hunched posture). Peripheral blood was routinely collected by tail vein incision into EDTA coated Microvette capillary blood collection tubes (Sarstedt AG & Co). Plasma was isolated after centrifugation (2000xg, 4°C, 4min) of blood samples and plasma IgG2b quantified by ELISA as described below. Animals were euthanized at ethical endpoint based on clinical symptoms and overall survival rates were assessed using Kaplan-Meier analysis.

For cross-sectional studies, bone marrow and spleen cells from mice were collected after 3 weeks of treatment as previously described (67). Bone marrow cells were isolated using a mortar and pestle and resuspended in FACS buffer. Spleens were weighed, pulverized to achieve single cells and resuspended in FACS buffer. Cells were filtered using a 70μM filter and red cells lysed using ACK lysis buffer (150mM NH_4_Cl, 10mM KHCO_3_, 0.1mM EDTA). Cells were then washed in FACS buffer and stained for flow-cytometry and FACS.

### Enzyme-linked ImmunoSorbent Assay (ELISA)

Assays were conducted as previously described(68). 96-well U-bottom plate (Corning, #CLS3795) were incubated with 50µL/well of 2 µg/mL goat anti-mouse IgG2b (southern Biotech #1090-01) for 5h at room temperature in humid conditions. Plates were washed with PBS/tween (500mL PBS + 3 drops of Tween20), PBS then distilled water, each three times. Plasma samples were serially diluted in blocking buffer (PBS, 1% FCS, 0.05% Tween20, 0.6% skim milk powder) and then titrated onto the pre-coated plates in blocking buffer along with the mouse myeloma protein IgG2b (Kappa) standard (MP Biomedicals #0850331). Plates were incubated overnight at room temperature in humid conditions, then washed as previously described. Goat anti-mouse IgG2b HRPO (Southern Biotech #1090-05) diluted 1:1000 in blocking buffer was transferred to the plates (50µL/well) and incubated at room temperature for 4h in humid conditions. Plates were washed as previously described and 100µL of developing solution (0.1M citric acid, 0.54 mg/mL ABTS, H_2_O_2_ 1X, ddH2O) added to each well. Plates were incubated for 30min in the dark prior to analyzing on the Cytation C10 Confocal Imaging Reader (BioTek) using wavelengths 415 minus 492.

### Flow cytometry and cell sorting

Cells were analyzed on the BD LSR Fortessa X-20 or BD FACSymphony or sorted on the BD FACSAria Fusion 5 or BD FACSAria Fusion 3 (BD Biosciences) at the Peter MacCallum Cancer Centre flow cytometry core facility. Data was analyzed using FlowJo (v 10.4). Cells were suspended in FACS buffer (2% FBS in PBS) and stained with fluorophore-conjugated antibodies where indicated.

### Competitive proliferation assays

Cell lines expressing Cas9 were transduced with sgRNA constructs in the FgH1tUTG-GFP (Addgene plasmid #70183) backbone and sgRNA-expressing cells (GFP^+^) were sorted by FACS. GFP^+^ cells were mixed in a 1:1 ratio with untransduced cells (GFP-) or BFP^+^ cells expressing a Scrambled sgRNA in a FgH1tUT-BFP backbone. Co-cultures were treated with doxycycline (1µg/ml) for 3 days to induce sgRNA expression prior to beginning the assay. Cells were treated with vehicle (DMSO) or VTP-50469 (200nm or 400nM) on Day 0 of the assay. Every 3-4 days, cells were passaged, drugs refreshed, and the ratio of GFP^+^ to non-GFP^+^ cells was analyzed by flow cytometry and normalized to the ratio at Day 0.

### Proliferation and viability assays

RPMI and MM1.S cells transduced with doxycycline-inducible sgRNAs guides targeting MEN1 (or control SCR) were plated at 0.2-0.3x10^6^/mL in triplicate and treated with doxycycline (1 µg/mL) for 3 days before beginning the assay. Cells were counted with Trypan Blue and passaged every 3-4 days and doxycycline replenished on Day 3.

For iMenin growth assays, MM cells were seeded at ∼0.1 x 10^6^/mL in a 96-well U-bottom plate in quadruplicate and treated with DMSO or VTP50469 (500 nM) (MedChemExpress #MCE-HY-114162) for a period of 10 days. Viable cells were quantified by flow cytometry with DAPI (Merck #D9542-10mg) on days 3, 5, 7 and 10 post-treatments. Cells were passaged at each time point where drug was also refreshed. The growth index was calculated using the GR method. To predict the growth index, we first used the 10 cell lines with both iMenin growth index and dependency data to fit a linear model of growth index as a function of MEN1 dependence (Figure S1E). To predict the growth index of the cell lines that were absent from the panel, we randomly drew from a normal distribution centred on the linear model with a standard deviation equal to the residual standard error of the linear model.

Cells were seeded at 2 x 10^5^ cells/mL (JJN-3), 3 x 10^5^ cells/mL (TK26, TK27, U266, OPM2, KMS-34), or 4 x 10^5^ cells/mL (KMS-11, MM1s) in triplicate in 96-well plates. Cells were treated with Vehicle (DMSO) or VTP-50469 (500nM), and/or Pomalidomide (400nM) or Dexamethasone (20nM) as indicated for 7 days. Cells were passaged and drug refreshed every 2-3 days. On days 4 and 7, 50 µL of cells were transferred to a new plate, stained with DAPI (1 µg/mL), and quantified by flow cytometry using DAPI exclusion.

Synergy assays using P300/CBP and Menin inhibitors were performed by seeding MM cells at ∼0.1 x 10^6^/mL in 96-well U-bottom plates and treating with VTP-50469 and A485 (cat#) or GNE781 (cat#) in a checkerboard format as indicated. Two days after treatment, cells were passaged and drug refreshed. Cell viability was assessed on day 4 by flow cytometry with DAPI (Merck #D9542-10mg).

Primary myeloma samples were seeded at 2.0 × 10^5^ cells/mL and treated with menin inhibitor (500 nM) alone or in combination with CBP inhibitor (960 nM) for 72 h. Cells were then stained with CD45-FITC and CD38-PerCP antibodies (BD Biosciences) for 15 min at room temperature, washed with FACS buffer (0.5% FCS in PBS), and fixed in 2% paraformaldehyde (PFA) on ice for 20 min. Following fixation, cells were washed and stained with Apo 2.7-PE (Immunotech Beckman Coulter) in permeabilisation buffer (0.3% saponin and 1% FCS in PBS) for 20 min on ice. After a final wash in FACS buffer, cells were resuspended in 300 μL FACS buffer. Samples were acquired using a Beckman Coulter CytoFLEX flow cytometer and analysed with FlowLogic software.

### Western Blotting

Cells were seeded at 0.2-0.3 x 10^6^/mL and treated with drugs as indicated or 1µg/mL doxycycline for 5 days to induce sgRNA expression. 1 x 10^6^ cells were lysed in Laemmli lysis buffer (60 mM Tris–HCl, 10% (v/v) Glycerol, 2% (w/v) SDS, 0.5 U benzonase nuclease, cOmplete mini protease inhibitor cocktail (Merck #5892953001)) for 30 min at 4°C. Protein concentration was determined using Pierce^TM^ BCA Protein Assay Kit as per manufacturer’s instructions (Thermo Scientific #A55865).

To confirm Menin knockout in sgMEN1 transduced cells, 4x reducing sample buffer (8% (w/v) SDS, 40% (v/v) glycerol, 10% (v/v) b-mercaptoethanol, 100mM Tris-HCl pH 6.8, 0.01% (w/v) Bromophenol Blue) was added and samples were incubated at 95°C for 10 min. Equal amounts of protein lysate were resolved on a 4-20% Mini-PROTEAN TGX Precast Protein Gel (Bio-Rad) and wet transferred onto an Immobilon-FL PVDF membrane 0.45 μm (Merck). The membrane was dried for 30 minutes at room temperature and blocked in Licor (TBS) Blocking Buffer at 4°C overnight. It was then incubated for 1 hour at room temperature in the following primary antibodies, diluted 1:5000 in Licor (TBS) Blocking Buffer + 0.2% Tween20: rabbit anti-Menin (A300-105A, Bethyl Laboratories), and mouse anti-β-Actin (A2228, Sigma-Aldrich). The membrane was then washed in TBS-T and incubated for 1 hour at room temperature in the following secondary antibodies, diluted 1:20000 in Licor (TBS) Blocking Buffer + 0.2% Tween20 + 0.01% SDS: IRDye 680RD goat anti-Mouse IgG (926-68070, LI-COR Biosciences), and IRDye 800CW goat anti-rabbit IgG (926-32211, LI-COR Biosciences). Then membrane was washed again in TBS-T followed by TBS, and imaged via near-infrared Western blot detection using the Odyssey CLx Imaging System and Image Studio software (LI-COR Biosciences).

For all other samples, 4x LDS sample buffer (NuPAGE^TM^ #NP0007), 10x sample reducing agent (NuPAGETM #NP0009) was added and samples were incubated at 70°C for 10 mins. Protein lysates were resolved on 4-12 % Bis-Tris Gel (NuPAGETM #NP0336BOX) and immunoblotted onto Nitrocellulose membrane 0.45 μm (BioRad #1620094). Membranes were incubated with primary and secondary antibodies, and near-infrared Western blot detection was performed by the Odyssey CLx imaging system (LI-COR Biosciences). The following antibodies were used: Menin rabbit polyclonal (1:10000)(Thermo Fisher #A300-105A), IRF-4 rabbit (1:1000)(Cell Signaling Technology #4964S), c-Myc rabbit (1:1000)(Cell Signaling Technology #9402), β-actin mouse (1:20000)(Sigma-Aldrich #A2228), IRDye 680RD goat anti-Mouse IgG (H + L) (1:20000)(LI-COR Biosciences #926-68070), IRDye 800CW goat anti-rabbit IgG (H + L) (1:40000)(LI-COR Biosciences #926-32211).

### Poolseq

3.0 x10^6^ JJN3 and RPMI cells were seeded at a density of 0.3 x 10^6^ cells/mL in triplicate, whereas 6.0 x10^6^ U266 cells were seeded at a density of 0.6 x 10^6^ cells/mL, into a T25 flask. Cells were cultured in DMSO and VTP-50469 (500nM) for the indicated timepoints, at which times 2.0 x10^6^ cells were lysed in Trizol reagent (Thermo Fisher) and RNA was extracted using the Direct-zol RNA Miniprep (Zymo Research), RNA was quantified using the Tapestation 4150. cDNA was synthesized with specific RT primers (Cat#) that allowed us to combine several samples into 4 pools. cDNA was amplified with Kapa HiFi HotStart Ready Mix (Cat#). Cleanup of adaptor ligated cDNA was performed using MagBIO beads.

Libraries were prepared using the 3′Pool-seq method as previously described(69). First-strand cDNA synthesis was performed by first mixing 200 ng of RNA (diluted to 5 μl with water) with 1 μl indexed reverse transcription (RT) primer (10 μM) and 1 μl dNTPs and incubating at 72 °C for 3 min. Then 10 μl of RT master mix (3.6 μl SuperScript 5× buffer, 0.25 μl water, 0.25 μl DTT 100 mM, 2 μl betaine 5 M, 0.9 μl MgCl_2_ 100 mM, 2.5 μl template switching oligo and 0.5 μl SuperScript II reverse transcriptase) was added to each sample and then incubated at 42 °C for 90 min followed by 10 cycles (50 °C for 2 min, 42 °C for 2 min) and 70 °C for 15 min. Samples with unique RT indexes were then pooled to a total volume of 20 μl for Exonuclease1 treatment by adding 1 μl of Exo1 and incubating at 37 °C for 45 min followed by 92 °C for 15 min. Exo1-treated pools were then cleaned using SPRI Select magnetic beads at a 0.6× ratio according to manufacturer’s instructions and eluted in 10 μl. Next, 15 μl of cDNA amplification master mix (12.5 μl KAPA HotStart Mix, 1.25 μl enrichment primer A 20 μM and 1.25 μl enrichment primer B 20 μM) was added to each pool before touch-up PCR: 95 °C for 3 min, 4 cycles (98 °C 20 s, 65 °C 45 s, 72 °C 3 min), 9 cycles (98 °C 20 s, 67 °C 20 s, 72 °C 3 min) and 72 °C for 5 min. Amplified cDNA was cleaned as before but eluted in 20 μl and then diluted to 0.25 ng μl^−1^. This was then subjected to tagmentation and PCR following the manufacturer’s instructions with the following changes: 2 μl of diluted cDNA was added to 4 μl of TD buffer followed by 2 μl of ATM buffer. For the amplification, a master mix was prepared (2 μl indexed i5 primer 2 μM, 2 μl enrichment primer A 2 μM and 6 μl NPM) and added to each tagmented pool. PCR reaction was carried out with 13 cycles. The pools were then cleaned as before and then run on a D5000 Tapestation tape for pooling before sequencing on the Illumina NovaSeq 6000. Samples were sequenced 150bp paired-end to a depth of ∼10x10^6^ reads/sample.

### RNAseq

RPMI8226 and MM1.S cells were transduced with dox-inducible sgRNAs targeting MEN1 and KMT2A were seeded at 0.25x10^6^/mL in triplicate. RPMI8226 cells were treated with doxycycline (2µg/mL) for 3 days before lysis and RNA extraction. MM1.S cells were passaged at 3 days with doxycycline refreshed and harvested on day 5.

MM1S cells were seeded at a density of ∼0.2x10^6^cells/mL in triplicate and cultured in DMSO, VTP-50469 (400nM), GNE781 (60nM) or the combination for 3 days prior to lysis and RNA extraction.

VkMYC-32052 cells from the bone marrow of mice treated with Revumenib or vehicle for 3 weeks were isolated by FACS (CD138+).

1-2x10^6^ human MM cells and 50,000-100,000 VkMYC-32052 cells were lysed in Trizol reagent (Thermo Fisher) and RNA was extracted using the Direct-zol RNA Miniprep and Microprep kits (Zymo Research), respectively. RNAseq was performed at the Peter MacCallum Cancer Centre Molecular Genomics core facility. RNAseq libraries were generated using the QuantSeq 3’ mRNA-seq Library Prep Kit for Illumina (Lexogen) and sequenced on an Illumina NextSeq500 to produce 100bp single-end reads with a depth of 3-10x10^6^ reads per sample.

### Transcriptomic analysis

Sequencing reads were de-multiplexed using bcl2fastq (v2.17.1.14), low quality reads (Q < 30) removed and trimmed using cutadapt (v1.14). VkMYC RNAseq reads were mapped to the mouse (mm10) reference genome using or STAR (v 2.5.3a). RNAseq in human cell lines were mapped to human (hg19) reference genomes using HISAT2 (v2.0.4). Reads from Poolseq samples were mapped to human (hg38) reference genome using or STAR (v 2.5.3a). Reads were counted using featureCounts from subread (v 2.0.0). Immunoglobulin transcripts were removed then counts were filtered using filterByExpr from edgeR (4.6.3) with default parameters, quantile normalized using voom and differential gene expression performed using Limma (v3.64.1). Raw RNAseq counts in RPMI8226 and MM1S IRF4 knockout cells were downloaded from GEO (GSE148984). RNAseq in VTP-50469 treated AML cells were downloaded from the supplementary tables(27,41). Gene set enrichment analysis was performed using fgsea (v.1.34.2) using the MSigDB (v2026.1) gene sets. Figures were generated in R (v 4.5.0) using ggplot2 (v 3.5.2), pheatmap (v1.0.13) and ComplexHeatmap (v2.24.1). Limma (v3.64.1) was used to generate barcodeplots and perform rotating gene set testing.

### ChIPseq

RPMI8226, SKMM2 and MM1S cells were seeded at 0.2-0.5x10^6^/mL and treated with DMSO, VTP-50469, GNE781 or the combination as indicated for 3 days. Samples were fixed with formaldehyde solution (1% w/v) for 15mins with gentle rocking and quenched by adding 2.5M glycine for 5mins. 2x10^7^ cells were washed twice in ice cold PBS and resuspended in 1ml of ChIP Lysis buffer (1% SDS (w/v), 10mM EDTA, 50mM Tris-HCl pH 8.0, cOmplete mini EDTA-free protease inhibitor cocktail (Sigma-Aldrich)). SKMM2 samples were sonicated using a Covaris ME220 Focused Ultrasonicator for 20 minutes Peak Power: 75.0, Duty Factor: 15.0 %, Average Power: 11.3, Temperature Set Point: 8 degrees C, Cycles/Burst: 1000). MM1.S samples were sonicated using a Covaris S220 Focused Ultrasonicator for 40 min (30s on (5% duty cycle; intensity - 5200 cycles per burst), 20s off). 9ml of modified RIPA buffer [1% (v/v) TritonX100, 0.1% (v/v) deoxycholate, 90mM NaCl, 10mM Tris-HCl pH8, cOmplete mini EDTA-free protease inhibitor cocktail (Sigma-Aldrich)] was added to each sample. 5% from each sample was collected and pooled for an input sample. 5.5µg of Menin (Thermo Fisher #A300-105A), MLL1 (Bethyl Laboratories #A300-086A), H3K4me3 (Abcam #ab8580) and H3K27ac (Abcam #ab4729) antibodies were added to indicated samples. Samples were rotated overnight at 4°C. Protein A Dynabeads (Theromo Fisher) were added to samples and incubated at 4°C for 2-4h. Dynabeads were washed twice in low salt wash buffer (0.1% (w/v) SDS, 1% (v/v) TritonX100 v/v, 2mM EDTA, 150mM NaCl, 20mM Tris-HCl pH 8), once in high salt buffer (0.1% (w/v) SDS, 1% (v/v) TritonX100 v/v, 2mM EDTA, 500mM NaCl, 20mM Tris-HCl pH 8) and once in TE buffer. Samples were eluted in reverse crosslinking buffer (1% (w/v) SDS, 100mM NaHCO_3_) and incubated at 65°C in a thermal mixer at 1000 rpm overnight. DNA was extracted with Min-Elute PCR Purification kit (Qiagen #28004). Sequencing libraries were prepared using NEBNext Ultra II DNA Library Prep Kit for Illumina kit (New England Biolabs) according to the manufacturer’s protocol. Cleanup of adaptor ligated DNA was performed using AMPure XP beads. Purified DNA was amplified using PCR enrichment with 12 and 16 cycles for input and ChIP samples respectively.

The ChIPseq experiment in RPMI8226 cells was sequenced on an Illumina NovaSeq6000 with 120bp paired-end reads to a depth of ∼4-10 million reads per sample. The Menin ChIPseq libraries in MM1S and SKMM2 cells were sequenced on an Illumina Nextseq2000 with 100bp single-end reads to a depth of 10-15 million reads per sample. The libraries from the ChIPseq experiment in MM1S cells treated with VTP-50469 and/or GNE781 were sequenced on an Illumina NovaSeq with 120bp single-end with ∼10 million reads per sample. However, fewer than 1 million reads were obtained for the input, therefore the input sample for this experiment was re-sequenced on the Illumina NovaSeq6000 to obtain greater coverage (∼9 million reads with 150bp paired-end sequencing). For down-stream analyses, only read 1 was used for the input and treated as single-end sequenced.

Raw sequencing reads were demultiplexed using bcl2fastq (v2.17.1.14) and adaptor sequences were removed using Trim Galore (-q 30) (v0.4.4) and reads aligned to the human genome (hg38) using Bowtie2 (v2.3.3) using default parameters. Samtools (v1.9) was used for processing SAM and BAM files. Picard (v2.6.0) was used to remove duplicate reads. BAM files were converted to BigWig files using deeptools (v3.5.0) bamCoverage (-bl $blacklist --effectiveGenomeSize 2913022398--binSize 10 --smoothLength 50 --normalizeUsing RPKM). H3K27Ac peaks were called using csaw (v1.42.0). Read parameters were defined with readParam with minq=20, reads in blacklisted regions discarded and reads restricted to autosomal or X chromosomes. For each sample, sliding windows (width=1000) were tiled across the genome at intervals of width/2. Windows were filtered relative to input using the filterWindowsControl function where the IP enrichment was 3-fold greater than input. Additionally, lowly abundant windows were discarded if log_2_CPM < 1. Nearby windows were merged using the mergeWindows function with a maximum width of 50000bp and maximum distance between adjacent windows of 1000bp, then annotated to nearest gene and genomic feature using ChIPSeeker.

Super-enhancers (SEs) were identified by calling peaks with macs2 (v2.1.1; --nomodel --extsize 100), discarding blacklisted regions with bedtools intersect (-v) and stitching together constituent enhancers that occurred within 12.5kb, excluding those that were within 2kb of a TSS, with ROSE2 (v1.0.5). H3K27Ac non-promoter peaks (>2kb from TSS) that did not overlap with SEs were classified as regular enhancers. Menin abundance at SEs was determined using deeptools multiBigwigSummary using the RPKM normalised bigwig files. Menin was considered bound to SEs if the log2FC between RPMI8226 DMSO Menin ChIP and input was greater than 0.7, lowly bound when 0.25<logFC≤0.7 and not bound when logFC≤0.25. Since there was a high correlation between the publicly available H3K27Ac HiChIP data in RPMI8226 cells (GSE267004)(42) and our H3K27Ac ChIPseq in RPMI8226 cells (r = 0.75 using multiBamSummary from deeptools v3.5.3), the genes regulated by SEs were annotated by overlapping the HiChIP data with our SEs and limiting the number of SE-to-promoter interactions to the top 2 most highly expressing genes at baseline. MM1S SEs were annotated to nearest gene using ChIPSeeker (v1.44.0) since the correlation between our H3K27Ac ChIPseq in MM1S cells and the H3K27Ac HiChIP data (42) in MM1S cells was weak (r = 0.5).

Publicly available ChIPseq datasets were downloaded from GEO (GSE49224, GSE246436) using SRA-Toolkit (v24.7.1) then trimmed and mapped as described above.

Deeptools multiBigwigSummary was used to quantify the signal over the defined genomic regions and ComplexHeatmap (v2.24.1) or deeptools (v3.5.3) were used to generate heatmaps and average signal summary plots. trackViewer (v1.44.0) was used to generate figures of ChIPseq signal tracks. clusterProfiler (v4.16.0) was used for gene ontology analysis of Menin-bound, VTP-50469 down-regulated genes.

### RIME

25 x 10^6^ RPMI8226 cells were plated at 0.5 x 10^6^/mL per replicate (n = 4), where 40 x 10^6^ were harvested two days later. Cells were fixed in 1% w/v Formaldehyde (Thermo Scientific #410731000) for 10 minutes with gentle rocking and quenched with 2.5M Glycine (0.125M final concentration) for 5 minutes, washed with ice cold PBS twice, pelleted and frozen at - 80°C in Protein LoBind tubes (Eppendorf #0030108116) until the next day. Simultaneously, Dynabeads^TM^ Protein A (Thermo Fisher Scientific #10-002-D) were added to fresh LoBind tubes, placed on a magnet (Sergei Lab Supplies #1005) washed four times with Pierce^TM^ Protein-Free Blocking Buffer (Thermo Scientific #23228), and incubated with 5ug of either IgG rabbit isotype control (Thermo Fisher #03-610-2) or Menin polyclonal (Thermo Fisher #A300-105A) antibodies rotating at 4°C overnight. The next day, fixed pellets were thawed on ice and lysed sequentially with three lysis buffers containing 1x cOmplete mini protease inhibitor cocktail (PI) (Merck): LB1 (50 mM HEPES pH 7.5, 140 mM NaCl, 1 mM EDTA, 10 % Glycerol, 0.5 % NP-40, 0.25% Triton X-100 in ddH_2_O), LB2 (10 mM Tris-HCl pH 8, 200 mM NaCl, 1 mM EDTA, 0.5 mM EGTA in ddH_2_O) and LB3 (10 mM Tris-HCl pH 8, 100 mM NaCl, 1 mM EDTA, 0.5 mM EGTA, 0.1 % Na-Deoxycholate, 0.5 % N-lauroylsarcosine in ddH_2_O). After adding LB3, samples were transferred into milliTUBE 1 mL AFA Fiber tubes (COVARIS #520130) then sonicated using the ME220 COVARIS sonicator for 25 minutes (Peak Power: 75.0, Duty Factor 15%, Cycles/Burst: 1000, Average Power: 11.3, Temperature Set Point: 8°C). Lysates were placed into LoBind tubes with 1% Triton X-100 added and centrifuged at 20,000 g for 10 minutes at 4°C. Clarified supernatant was placed into new LoBind tubes, followed by the addition of RNAse A (Cat#) for 10 minutes at 37°C. Antibody-incubated beads were washed three times with blocking buffer and wash in LB3 containing 1x PI with 1% Triton X-100. After washing, pre-cleared lysates were equally divided into their respective IgG and Menin antibody-bead-containing LoBind tubes and incubate rotating at 4°C overnight. The following day, tubes were placed on a magnetic rack, supernatant discarded and beads were washed ten times with wash buffer (50mM HEPES pH 7.6, 1mM EDTA, 0.7% Na deoxycholate, 1% NP-40, 0.5M LiCL), then eluted in urea-SDS lysis/solubilization buffer (5% SDS, 8M urea, 10 mM glycine pH 7.55) and samples were frozen in –80°C overnight. Samples were purified using the S-Trap^TM^ Micro High Recovery protocol and columns (Cat#), with a few modifications to the steps: MMTS was replaced with 500 mM Iodoacetamide (generous gift from the Johnstone lab) and incubated at 30 minutes, omitted the addition of 0.2% formic acid in the final elution step and SpeedVac of the samples to dry and pellet proteins in regular tubes. Samples were placed in -80°C until they were ready to be submitted for sequence. Samples were resuspended in reconstitution buffer (2% acetonitrile, 0.5% TFA), vortexed, sonicated, centrifuged and transferred into mass spectrometry grade sample vials (Thermo #11190933).

LC-MS/MS was carried out using a Orbitrap Ascend mass spectrometer (Thermo Scientific) equipped with a nanoflow reversed-phase-HPLC (Ultimate 3000 RSLC, Dionex) fitted with an Acclaim Pepmap nano-trap column (Dionex—C18, 100 Å, 75 µm× 2 cm) and an Acclaim Pepmap RSLC analytical column (Dionex—C18, 100 Å, 75 µm× 50 cm). The tryptic peptides were injected to the enrichment column at an isocratic flow of 5 µL/min of 2% v/v CH_3_CN containing 0.1% v/v formic acid for 5 min applied before the enrichment column was switched in-line with the analytical column. The eluents were 5% DMSO in 0.1% v/v formic acid (solvent A) and 5% DMSO in 100% v/v CH_3_CN and 0.1% v/v formic acid (solvent B). The flow gradient was (i) 0-6min at 3% B, ii) 6-7min, 3-4% (ii) 7-82 min, 4-25% B (iii) 82-86min 25-40% B (iv) 86-87min, 40-80% B (v) 87-90min, 80-80% B (vi) 90-91min, 80-3% and equilibrated at 3% B for 10 minutes before the next sample injection.

For DIA experiments full MS resolutions were set to 120,000 at m/z 200 and scanning from 350-1400m/z in the profile mode. Full MS AGC target was 250% with an IT of 50 ms. AGC target value for fragment spectra was set at 2000%. 50 windows of 13.7 Da were used with an overlap of 1 Da. Resolution was set to 30,000 and maximum IT to 55 ms. Normalized collision energy was set at 30%. All data were acquired in centroid mode using positive polarity.

Raw files were searched using the direct DIA analysis workflow with default settings on the Spectronaut software (v. 20.2.250922.92449) and against the reviewed Uniport *Homo Sapiens* database (download Nov 2025). For Spectronaut searches, Trypsin specificity was set to two missed cleavages. Carbamidomethyl (Cys) was defined as fixed modification and acetylation (protein N-term) and oxidation (Met) as variable modification. Search results are filtered at a protein, peptide and PSM false discovery rate of 1%. DIA analysis identification had a PEP and QValue cutoff at 0.01. Precursor filtering using the Qvalue, quantification carried out on the MS2 level and cross-run normalization strategy set to automatic. PG.Quantity output was used for relative quantitation and statistical analysis. Differential analysis was performed using DIA-Analyst (https://analyst-suites.org/apps/dia-analyst/) with missing not at random MinProb imputation.

## Notes

### Competing Interest Statement

The authors have declared no competing interest.

## References

1. Mafra A, Laversanne M, Marcos-Gragera R, Chaves HVS, Mcshane C, Bray F, et al. The global multiple myeloma incidence and mortality burden in 2022 and predictions for 2045. J Natl Cancer Inst. Oxford University Press (OUP); 2025;117:907–14.

2. Malard F, Neri P, Bahlis NJ, Terpos E, Moukalled N, Hungria VTM, et al. Multiple myeloma. Nat Rev Dis Primers. Springer Science and Business Media LLC; 2024;10:45.

3. Biltibo EMD, Surapaneni M, Al Hadidi S, Suvannasankha A, Jayani-Kosarzycki RV. Current and future directions of immunotherapies in multiple myeloma. Am Soc Clin Oncol Educ Book. 2025;45:e473316.

4. Whyte WA, Orlando DA, Hnisz D, Abraham BJ, Lin CY, Kagey MH, et al. Master Transcription Factors and Mediator Establish Super-Enhancers at Key Cell Identity Genes. Cell. Elsevier Inc.; 2013;153:307–19.

5. Hnisz D, Abraham BJ, Lee TI, Lau A, Saint-André V, Sigova AA, et al. Super-Enhancers in the Control of Cell Identity and Disease. Cell. Elsevier Inc.; 2013;155:934–47.

6. Bergsagel PL, Chesi M, Nardini E, Brents LA, Kirby SL, Kuehl WM. Promiscuous translocations into immunoglobulin heavy chain switch regions in multiple myeloma. Proc Natl Acad Sci U S A. Proceedings of the National Academy of Sciences; 1996;93:13931–6.

7. Chesi M, Nardini E, Brents LA, Schröck E, Ried T, Kuehl WM, et al. Frequent translocation t(4;14)(p16.3;q32.3) in multiple myeloma is associated with increased expression and activating mutations of fibroblast growth factor receptor 3. Nat Genet. Springer Science and Business Media LLC; 1997;16:260–4.

8. Chesi M, Bergsagel PL, Brents LA, Smith CM, Gerhard DS, Kuehl WM. Dysregulation of cyclin D1 by translocation into an IgH gamma switch region in two multiple myeloma cell lines. Blood. Blood; 1996;88:674–81.

9. Affer M, Chesi M, Chen W-DG, Keats JJ, Demchenko YN, Roschke AV, et al. Promiscuous MYC locus rearrangements hijack enhancers but mostly super-enhancers to dysregulate MYC expression in multiple myeloma. Leukemia. Springer Science and Business Media LLC; 2014;28:1725–35.

10. Maura F, Bolli N, Angelopoulos N, Dawson KJ, Leongamornlert D, Martincorena I, et al. Genomic landscape and chronological reconstruction of driver events in multiple myeloma. Nat Commun. 2019;10:3835.

11. Manier S, Salem KZ, Park J, Landau DA, Getz G, Ghobrial IM. Genomic complexity of multiple myeloma and its clinical implications. Nat Rev Clin Oncol. 2017;14:100–13.

12. Jin Y, Chen K, De Paepe A, Hellqvist E, Krstic AD, Metang L, et al. Active enhancer and chromatin accessibility landscapes chart the regulatory network of primary multiple myeloma. Blood. 2018;131:2138–50.

13. Ordoñez R, Kulis M, Russiñol N, Chapaprieta V, Carrasco-Leon A, García-Torre B, et al. Chromatin activation as a unifying principle underlying pathogenic mechanisms in multiple myeloma. Genome Res. Cold Spring Harbor Laboratory; 2020;30:1217–27.

14. Alvarez-Benayas J, Trasanidis N, Katsarou A, Ponnusamy K, Chaidos A, May PC, et al. Chromatin-based, in cis and in trans regulatory rewiring underpins distinct oncogenic transcriptomes in multiple myeloma. Nat Commun. 2021;12:5450.

15. Lovén J, Hoke HA, Lin CY, Lau A, Orlando DA, Vakoc CR, et al. Selective inhibition of tumor oncogenes by disruption of super-enhancers. Cell. Elsevier Inc.; 2013;153:320–34.

16. de Matos Simoes R, Shirasaki R, Downey-Kopyscinski SL, Matthews GM, Barwick BG, Gupta VA, et al. Genome-scale functional genomics identify genes preferentially essential for multiple myeloma cells compared to other neoplasias. Nat Cancer. 2023;4:754–73.

17. Neri P, Barwick BG, Jung D, Patton JC, Maity R, Tagoug I, et al. ETV4-Dependent Transcriptional Plasticity Maintains MYC Expression and Results in IMiD Resistance in Multiple Myeloma. Blood Cancer Discov. 2024;5:56–73.

18. Welsh SJ, Barwick BG, Meermeier EW, Riggs DL, Shi CX, Zhu YX, et al. Transcriptional Heterogeneity Overcomes Super-Enhancer Disrupting Drug Combinations in Multiple Myeloma. Blood Cancer Discov. 2024;5:34–55.

19. Nicosia L, Spencer GJ, Brooks N, Amaral FMR, Basma NJ, Chadwick JA, et al. Therapeutic targeting of EP300/CBP by bromodomain inhibition in hematologic malignancies. Cancer Cell. Elsevier BV; 2023;41:2136–2153.e13.

20. Hogg SJ, Motorna O, Cluse LA, Johanson TM, Coughlan HD, Raviram R, et al. Targeting histone acetylation dynamics and oncogenic transcription by catalytic P300/CBP inhibition. Mol Cell. Elsevier Inc.; 2021;81:2183–2200.e13.

21. Matkar S, Thiel A, Hua X. Menin: a scaffold protein that controls gene expression and cell signaling. Trends Biochem Sci. Elsevier Ltd; 2013;38:394–402.

22. Brown MR, Soto-Feliciano YM. Menin: from molecular insights to clinical impact. Epigenomics. Informa UK Limited; 2025;17:489–505.

23. Yokoyama A, Somervaille TCP, Smith KS, Rozenblatt-Rosen O, Meyerson M, Cleary ML. The Menin Tumor Suppressor Protein Is an Essential Oncogenic Cofactor for MLL-Associated Leukemogenesis. Cell. Elsevier Inc.; 2005;123:207–18.

24. Yokoyama A, Cleary ML. Menin Critically Links MLL Proteins with LEDGF on Cancer-Associated Target Genes. Cancer Cell. Elsevier Inc.; 2008;14:36–46.

25. Kühn MWM, Song E, Feng Z, Sinha A, Chen C-W, Deshpande AJ, et al. Targeting Chromatin Regulators Inhibits Leukemogenic Gene Expression in NPM1 Mutant Leukemia. Cancer Discov. American Association for Cancer Research; 2016;6:1166–81.

26. Borkin D, He S, Miao H, Kempinska K, Pollock J, Chase J, et al. Pharmacologic Inhibition of the Menin-MLL Interaction Blocks Progression of MLL Leukemia In Vivo. Cancer Cell. Elsevier Inc.; 2015;27:589–602.

27. Krivtsov AV, Evans K, Gadrey JY, Eschle BK, Hatton C, Uckelmann HJ, et al. A Menin-MLL Inhibitor Induces Specific Chromatin Changes and Eradicates Disease in Models of MLL-Rearranged Leukemia. Cancer Cell. Elsevier Inc.; 2019;36:660–673.e11.

28. Issa GC, Aldoss I, DiPersio J, Cuglievan B, Stone R, Arellano M, et al. The menin inhibitor revumenib in KMT2A-rearranged or NPM1-mutant leukaemia. Nature. Springer Science and Business Media LLC; 2023;615:920–4.

29. Arellano ML, Thirman MJ, DiPersio JF, Heiblig M, Stein EM, Schuh AC, et al. Menin inhibition with revumenib for NPM1-mutated relapsed or refractory acute myeloid leukemia: the AUGMENT-101 study. Blood. American Society of Hematology; 2025;146:1065–77.

30. Issa GC, Aldoss I, Thirman MJ, DiPersio J, Arellano M, Blachly JS, et al. Menin inhibition with revumenib for KMT2A-rearranged relapsed or refractory acute leukemia (AUGMENT-101). J Clin Oncol. American Society of Clinical Oncology (ASCO); 2025;43:75–84.

31. Wang ES, Montesinos P, Foran J, Erba H, Rodríguez-Arbolí E, Fedorov K, et al. Ziftomenib in relapsed or refractory NPM1-mutated AML. J Clin Oncol. American Society of Clinical Oncology (ASCO); 2025;43:3381–90.

32. Grembecka J, He S, Shi A, Purohit T, Muntean AG, Sorenson RJ, et al. Menin-MLL inhibitors reverse oncogenic activity of MLL fusion proteins in leukemia. Nat Chem Biol. Springer Science and Business Media LLC; 2012;8:277–84.

33. Heikamp EB, Henrich JA, Perner F, Wong EM, Hatton C, Wen Y, et al. The Menin-MLL1 interaction is a molecular dependency in NUP98-rearranged AML. Blood [Internet]. American Society of Hematology; 2021; Available from: http://eutils.ncbi.nlm.nih.gov/entrez/eutils/elink.fcgi?dbfrom=pubmed&id=34582559&retmode=ref&cmd=prlinks

34. Barajas JM, Rasouli M, Umeda M, Hiltenbrand R, Abdelhamed S, Mohnani R, et al. Acute myeloid leukemias with UBTF tandem duplications are sensitive to menin inhibitors. Blood. 2024;143:619–30.

35. Arafeh R, Shibue T, Dempster JM, Hahn WC, Vazquez F. The present and future of the Cancer Dependency Map. Nat Rev Cancer. Springer Science and Business Media LLC; 2025;25:59–73.

36. Shaffer AL, Emre NC, Lamy L, Ngo VN, Wright G, Xiao W, et al. IRF4 addiction in multiple myeloma. Nature. 2008;454:226–31.

37. Olaisen C, Røst LM, Sharma A, Søgaard CK, Khong T, Berg S, et al. Multiple myeloma cells with increased proteasomal and ER stress are hypersensitive to ATX-101, an experimental peptide drug targeting PCNA. Cancers (Basel). MDPI AG; 2024;16:3963.

38. Clark NA, Hafner M, Kouril M, Williams EH, Muhlich JL, Pilarczyk M, et al. GRcalculator: an online tool for calculating and mining dose-response data. BMC Cancer. Springer Science and Business Media LLC; 2017;17:698.

39. Palumbo A, Chanan-Khan A, Weisel K, Nooka AK, Masszi T, Beksac M, et al. Daratumumab, bortezomib, and dexamethasone for multiple myeloma. N Engl J Med. Massachusetts Medical Society; 2016;375:754–66.

40. Fedele PL, Liao Y, Gong JN, Yao Y, van Delft MF, Low MSY, et al. The transcription factor IRF4 represses proapoptotic BMF and BIM to licence multiple myeloma survival. Leukemia. 2021;35:2114–8.

41. Uckelmann HJ, Kim SM, Wong EM, Armstrong SA. Therapeutic targeting of preleukemia cells in a mouse model of NPM1 mutant acutemyeloid leukemia. Science. American Association for the Advancement of Science; 2020;367:586–90.

42. Xiong S, Zhou J, Tan TK, Chung TH, Tan TZ, Toh SH, et al. Super enhancer acquisition drives expression of oncogenic PPP1R15B that regulates protein homeostasis in multiple myeloma. Nat Commun. 2024;15:6810.

43. Jia Y, Zhou J, Tan TK, Chung TH, Wong RWJ, Chooi JY, et al. Myeloma-specific superenhancers affect genes of biological and clinical relevance in myeloma. Blood Cancer J. 2021;11:32.

44. Mohammed H, D’Santos C, Serandour AA, Ali HR, Brown GD, Atkins A, et al. Endogenous purification reveals GREB1 as a key estrogen receptor regulatory factor. Cell Rep. 2013;3:342–9.

45. Huang J, Gurung B, Wan B, Matkar S, Veniaminova NA, Wan K, et al. The same pocket in menin binds both MLL and JUND but has opposite effects on transcription. Nature. Nature Publishing Group; 2012;482:542–6.

46. Li W, Xu H, Xiao T, Cong L, Love MI, Zhang F, et al. MAGeCK enables robust identification of essential genes from genome-scale CRISPR/Cas9 knockout screens. Genome Biol. Genome Biology; 2014;15:819–812.

47. Szklarczyk D, Kirsch R, Koutrouli M, Nastou K, Mehryary F, Hachilif R, et al. The STRING database in 2023: protein-protein association networks and functional enrichment analyses for any sequenced genome of interest. Nucleic Acids Res. Oxford University Press (OUP); 2023;51:D638–46.

48. Weinert BT, Narita T, Satpathy S, Srinivasan B, Hansen BK, Scholz C, et al. Time-Resolved Analysis Reveals Rapid Dynamics and Broad Scope of the CBP/p300 Acetylome. Cell. 2018;174:231–244 e12.

49. Mashtalir N, D’Avino AR, Michel BC, Luo J, Pan J, Otto JE, et al. Modular Organization and Assembly of SWI/SNF Family Chromatin Remodeling Complexes. Cell. Elsevier Inc.; 2018;175:1272–1288.e20.

50. Nakayama RT, Pulice JL, Valencia AM, McBride MJ, McKenzie ZM, Gillespie MA, et al. SMARCB1 is required for widespread BAF complex-mediated activation of enhancers and bivalent promoters. Nat Genet. Springer Science and Business Media LLC; 2017;49:1613–23.

51. Bolomsky A, Ceribelli M, Scheich S, Rinaldi K, Huang DW, Chakraborty P, et al. IRF4 requires ARID1A to establish plasma cell identity in multiple myeloma. Cancer Cell. 2024;42:1185–1201 e14.

52. Zhao Z, Shilatifard A. Enhancer and metabolic rewiring by KMT2C-COMPASS or KMT2D-COMPASS family loss in cancer creates druggable vulnerabilities. Nat Rev Cancer [Internet]. Springer Science and Business Media LLC; 2026 [cited 2026 May 12]; Available from: 10.1038/s41568-026-00919-x

53. Pawlyn C, Kaiser MF, Heuck C, Melchor L, Wardell CP, Murison A, et al. The spectrum and clinical impact of epigenetic modifier mutations in myeloma. Clin Cancer Res. American Association for Cancer Research (AACR); 2016;22:5783–94.

54. Soto-Feliciano YM, Sanchez-Rivera FJ, Perner F, Barrows DW, Kastenhuber ER, Ho YJ, et al. A Molecular Switch between Mammalian MLL Complexes Dictates Response to Menin-MLL Inhibition. Cancer Discov. 2023;13:146–69.

55. Borbolis F, Syntichaki P. Biological implications of decapping: beyond bulk mRNA decay. FEBS J. Wiley; 2022;289:1457–75.

56. Aubrey BJ, Cutler JA, Bourgeois W, Donovan KA, Gu S, Hatton C, et al. IKAROS and MENIN coordinate therapeutically actionable leukemogenic gene expression in MLL-r acute myeloid leukemia. Nat Cancer. 2022;3:595–613.

57. Bourgeois W, Cutler JA, Aubrey BJ, Wenge DV, Perner F, Martucci C, et al. Mezigdomide is effective alone and in combination with menin inhibition in preclinical models of KMT2A-r and NPM1c AML. Blood. 2024;143:1513–27.

58. Joseph N, Campbell V, Pawlyn C, Vogl D, Holstein S, Gooding S, et al. Randomized phase II dose optimization study of inobrodib (CCS1477), in combination with pomalidomide and dexamethasone in relapsed/refractory multiple myeloma (RRMM). Blood. American Society of Hematology; 2025;146:4035–4035.

59. Yadav B, Wennerberg K, Aittokallio T, Tang J. Searching for drug synergy in complex dose-response landscapes using an interaction potency model. Comput Struct Biotechnol J. American Association for the Advancement of Science (AAAS); 2015;13:504–13.

60. Chesi M, Matthews GM, Garbitt VM, Palmer SE, Shortt J, Lefebure M, et al. Drug response in a genetically engineered mouse model of multiple myeloma is predictive of clinical efficacy. Blood. American Society of Hematology; 2012;120:376–85.

61. Maura F, Coffey DG, Stein CK, Braggio E, Ziccheddu B, Sharik ME, et al. The genomic landscape of Vk*MYC myeloma highlights shared pathways of transformation between mice and humans. Nat Commun. 2024;15:3844.

62. Chesi M, Robbiani DF, Sebag M, Chng WJ, Affer M, Tiedemann R, et al. AID-dependent activation of a MYC transgene induces multiple myeloma in a conditional mouse model of post-germinal center malignancies. Cancer Cell. 2008;13:167–80.

63. Chandrasekharappa SC, Guru SC, Manickam P, Olufemi SE, Collins FS, Emmert-Buck MR, et al. Positional cloning of the gene for multiple endocrine neoplasia-type 1. Science. American Association for the Advancement of Science (AAAS); 1997;276:404–7.

64. Adriaanse FRS, Schneider P, Arentsen-Peters STCJM, Fonseca AMN da, Stutterheim J, Pieters R, et al. Distinct responses to menin inhibition and synergy with DOT1L inhibition in KMT2A-rearranged acute lymphoblastic and myeloid leukemia. Int J Mol Sci. MDPI AG; 2024;25:6020.

65. Sharlandjieva V, Chahrour C, Lassen FH, Hamley JC, Damianou A, Denny N, et al. Menin maintains enhancer-promoter interactions in a leukemia-specific manner [Internet]. bioRxiv. bioRxiv; 2026. Available from: 10.64898/2026.01.16.698179

66. DiNardo CD, Savona MR, Kishtagari A, Fathi AT, Bhalla KN, Agresta S, et al. Preliminary results from a phase 1 dose escalation study of FHD-286, a novel BRG1/BRM (SMARCA4/SMARCA2) inhibitor, administered as an oral monotherapy in patients with advanced hematologic malignancies. Blood. American Society of Hematology; 2023;142:4284–4284.

67. Gruber E, So J, Lewis AC, Franich R, Cole R, Martelotto LG, et al. Inhibition of mutant IDH1 promotes cycling of acute myeloid leukemia stem cells. Cell Rep. Elsevier BV; 2022;40:111182.

68. Kong IY, Trezise S, Light A, Todorovski I, Arnau GM, Gadipally S, et al. Epigenetic modulators of B cell fate identified through coupled phenotype-transcriptome analysis. Cell Death Differ. 2022;29:2519–30.

69. Neville D, Ferguson DT, Heikamp EB, Lai Z, Magor GW, Lam C, et al. DOT1L provides transcriptional memory through PRC1.1 antagonism. Nat Cell Biol. Springer Science and Business Media LLC; 2026;28:307–22.

