## Supplemental material for "Menin regulates oncogenic cell identity transcriptional networks in multiple myeloma"

**Supplemental tables**

Table S1. DEGs for MM cell lines treated with 500nM VTP-50469 for 3 or 5 days.

Table S2. DEGs induced by MEN1 or KMT2A knockout in MM cell lines.

Table S3. SEs in RPMI8226 cells.

Table S4. Output from DIA-analyst for Menin RIME in RPMI8226 cells.

Table S5. Raw counts and MAGeCK outputs for CRISPR screen in MM1S cells treated with VTP-50469.

Table S6. Patient sample characteristics.

Table S7. DEGs in MM1S cells treated with VTP-50469 (400nM), GNE781 (60nM) or the combination for 3 days.

Table S8. DEGs in VkMYC-32052 cells from the bone marrow of mice treated with revumenib and/or inobrodib for 21 days.

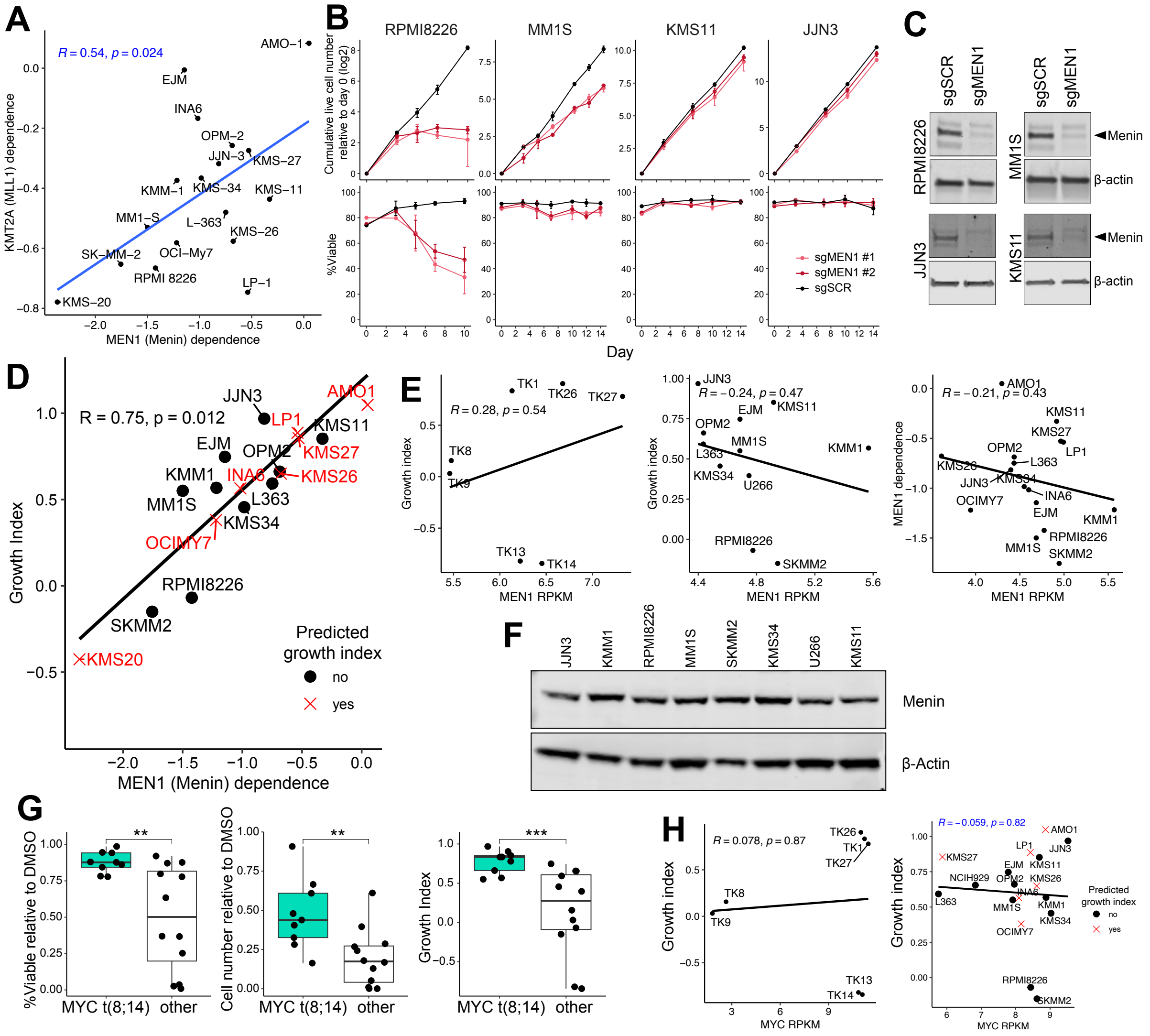


**Figure S1: Myeloma cells are sensitive to genetic and pharmacological disruption of Menin.**

**A.** Correlation between MEN1 and KMT2A DepMap dependency scores across myeloma cell lines.

**B.** Proliferation assays in MM cell lines transduced with sgRNAs targeting Menin. Cell number and viability (DAPI-) were assessed by flow-cytometry. Mean ± SD of 3 biological replicates.

**C.** Western blots in MM cell lines transduced with sgRNAs targeting Menin.

**D.** Correlation between MEN1 dependency scores (DepMap) and VTP-50469 growth index. Growth index was predicted using the linear model (see methods) for the red cell lines.

**E.** Correlation between menin RNA expression and VTP-50469 growth index in MM cell lines.

**F.** Menin protein expression across MM cell lines. Representative western blot from 2 biological replicates.

**G.** Viability and cell number for VTP-50469 treated MM cell lines stratified by the occurrence of the t(8;14) MYC translocation.

**H.** Correlation between MYC expression and VTP-50469 growth index.

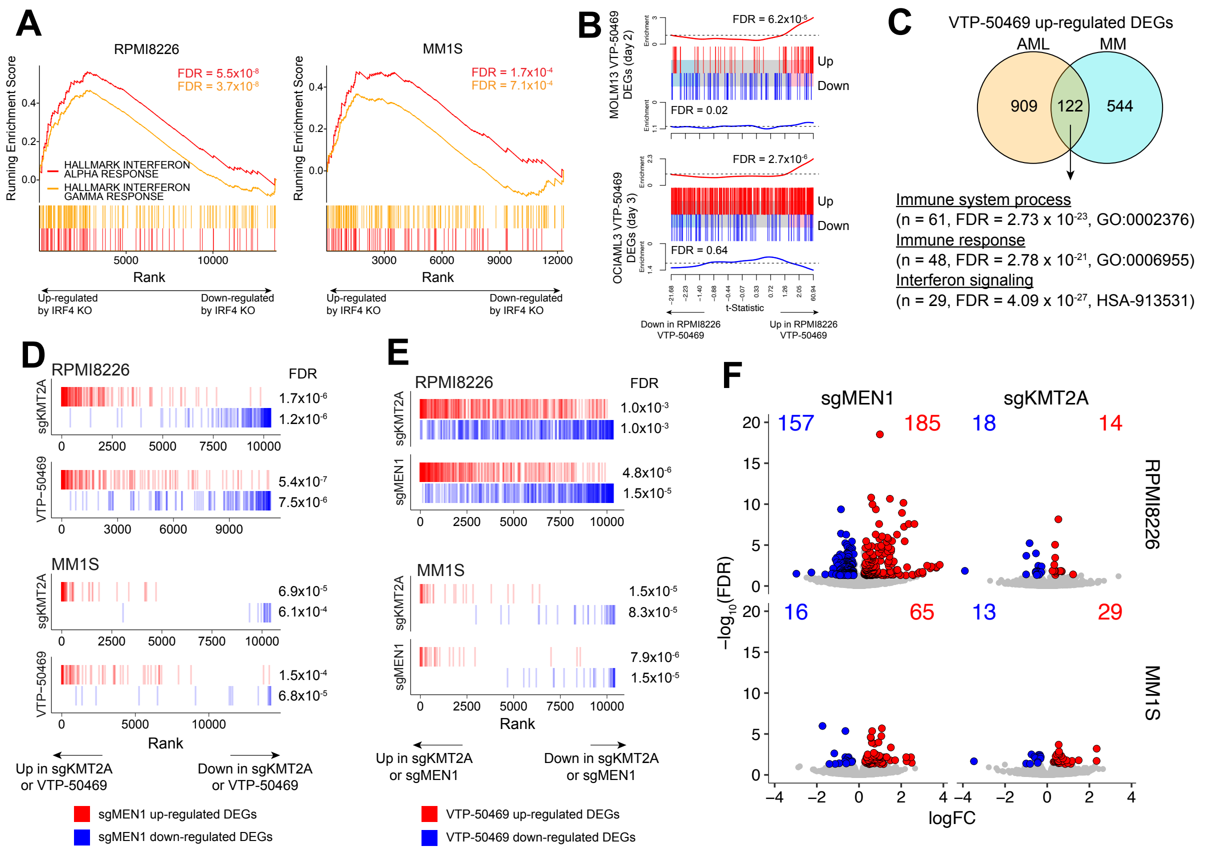


**Figure S2: Menin inhibition downregulates core myeloma regulatory networks.**

**A-F.** RNAseq in RPMI8226 or MM1S cells treated with DMSO or VTP-50469 (500nM and 400nM, respectively) for 3 days, or transduced with sgRNAs targeting MEN1 or KMT2A.

**A.** Ranked enrichment plot of interferon gene signatures.

**B.** Barcodeplot using genesets comprised of DEGs from AML cells treated with VTP-50469[1,2].

**C.** Venn diagram showing the overlap of VTP-50469 up-regulated DEGs across AML and MM models are enriched for immune pathways by STRING analysis.

**D-E.** Ranked enrichment plot comparing MEN1 knockout DEGs (D) and VTP-50469 DEGs (E) across given conditions. Significance assessed using ROAST.

**F.** Volcano plots highlighting DEGs.

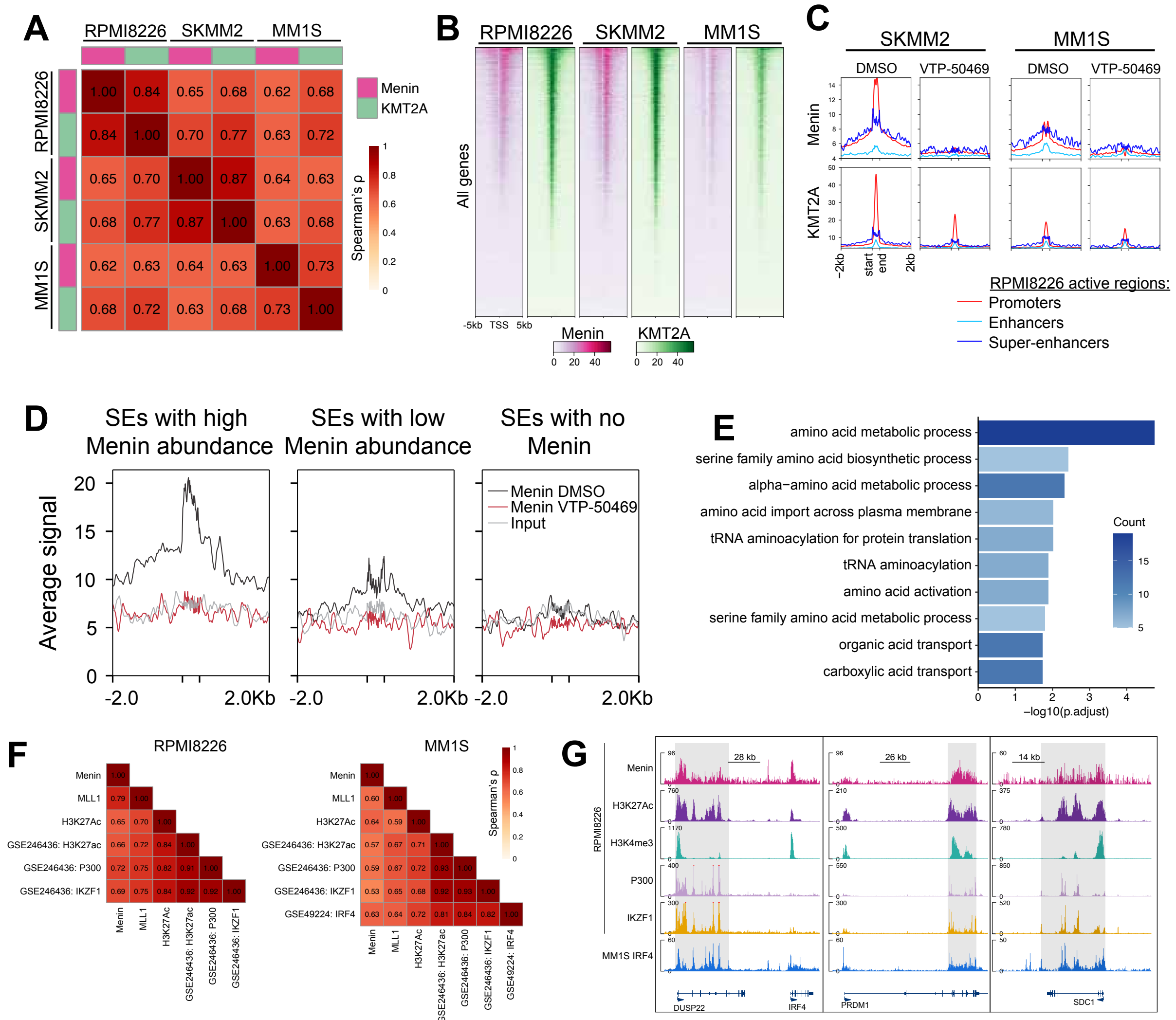


**Figure S3: Menin/KMT2A complex is present at promoters and SEs and is evicted from chromatin by VTP-50469.**

**A-E.** ChIPseq in RPMI8226, SKMM2 and MM1S cells treated with VTP-50469 (500nM) for 3 days.

**A.** Correlation between Menin and MLL1 ChIPseq tracks.

**B.** Tornado plot of Menin and MLL1 across the TSS of all genes.

**C.** Average Menin and MLL1 signal across genomic regions defined in RPMI8226 cells.

**D.** Average Menin signal at SEs stratified by Menin abundance.

**E.** Gene Ontology enrichment for Menin-regulated genes.

**F.** Correlation of ChIPseq tracks.

**G.** ChIPseq signal tracks. Shaded region denotes SEs in RPMI8226 cells.

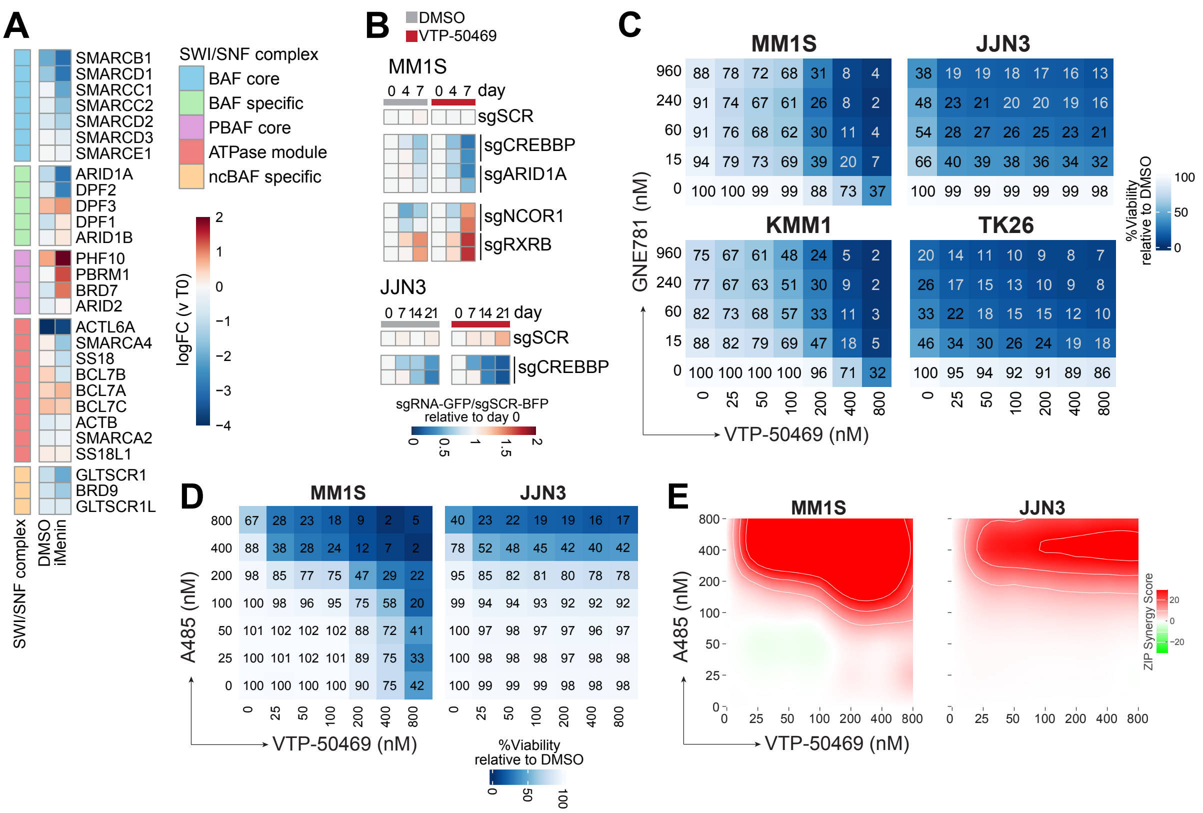

**Figure S4: EP300/NCOR1 axis is a key regulator of iMenin sensitivity.**

**A.** CRISPR screen in MM1S cells cultured for 32 days in increasing concentrations of VTP-50469 (200-400nM). Heatmap showing the average logFC for the sgRNAs targeting components of the SWI/SNF complexes.

**B.** Competition assays of myeloma cells transduced with sgRNA-GFP targeting given genes mixed 50:50 with control sgSCR-BFP cells. Cells were treated with 400nM VTP-50469 or DMSO.

**C-E.** Myeloma cells were treated with increasing concentrations of VTP-50469 and/or iEP300/CREBBP (GNE-781 or A485) for 4 days and viability assessed by DAPI staining and flow-cytometry. N=3 biological replicates. Average percent viable (DAPI-) (C-D) and ZIP synergy scores (E).

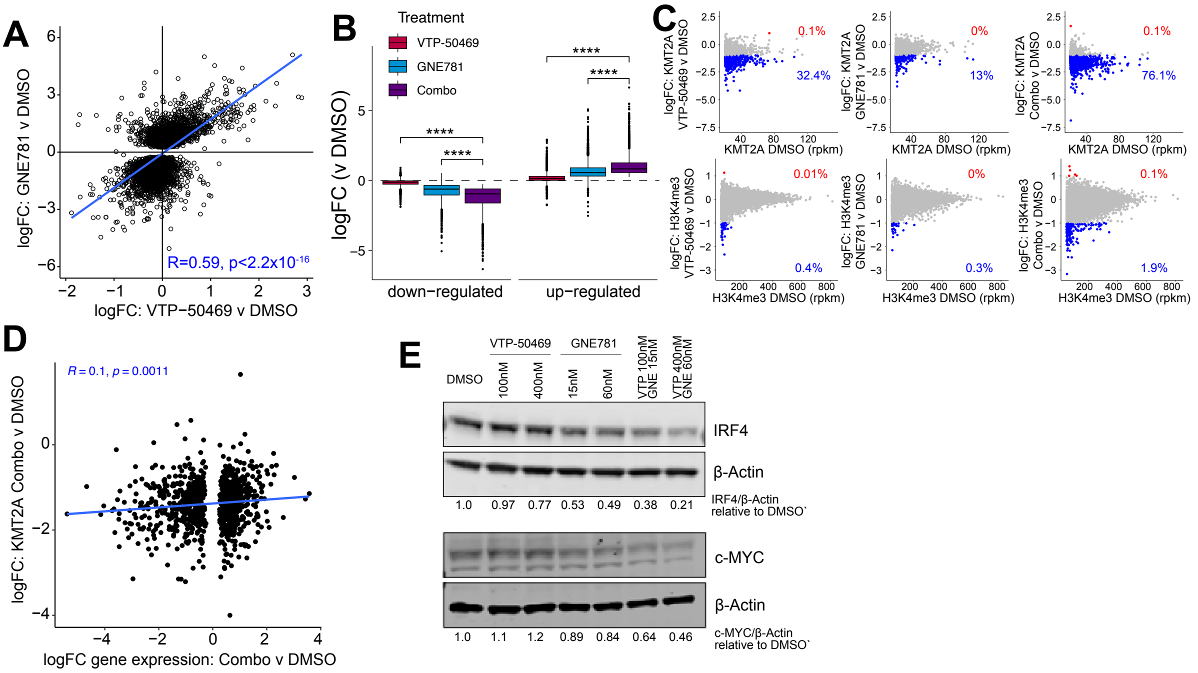


**Figure S5: iMenin and iEP300 synergize by suppressing SE networks.**

**A-D.** RNAseq and ChIPseq of MM1S cells treated with VTP-50469 (400nM), GNE-781 (60nM) or the combination for 3 days.

**A.** Correlation between single-agent VTP-50469 and GNE-781 DEGs (FDR < 0.05 and logFC < -0.25).

**B.** Boxplot of combination down DEGs (FDR < 0.05 and logFC < -0.25) across all conditions. Significance assessed with Student's t-test. ****p<0.0001.

**C.** Changes to KMT2A and H3K4Me3 at promoters across the different conditions.

**D.** Correlation between gene expression changes and changes in KMT2A at the promoters of DEGs under combination treatment.

**E.** Western blot in JJN3 cells treated with VTP-50469 and/or GNE-781 for 3 days.

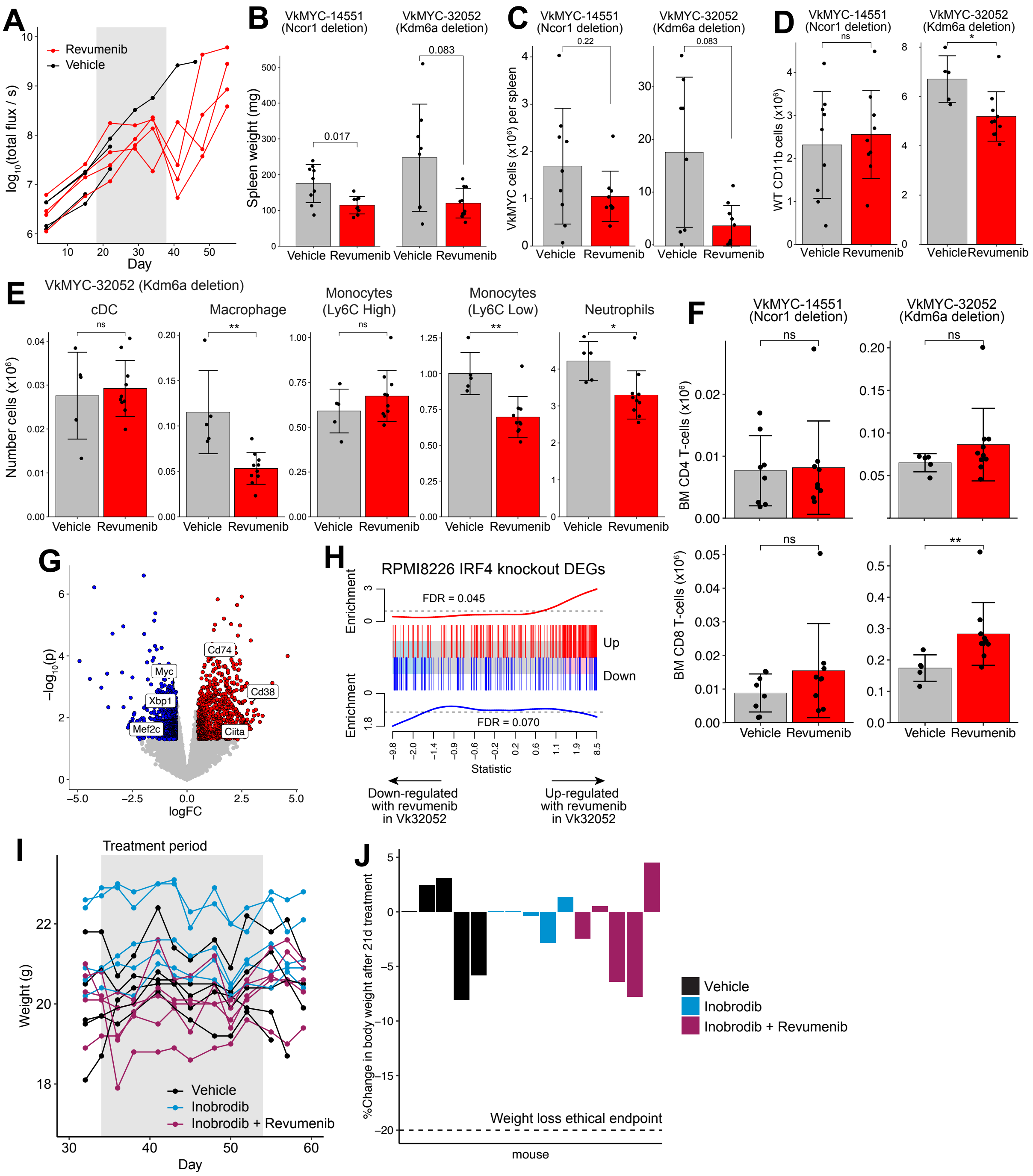

**Figure S6: Revumenib demonstrates anti-myeloma efficacy *in vivo* as a single agent and in combination with inobrodib.**

**A.** RPMI8226 cells expressing luciferase were transplanted by intrafemoral injection into NSG mice and treated with 50mg/kg Revumenib by oral gavage b.i.d. for 3 weeks. Total flux over time for each mouse.

**B-F.** VkMYC cells with a monoallelic *Ncor1* deletion (#14551) or biallelic *Kdm6a* deletion (#32052) were transplanted into C57BL/6 mice by tail vein injection and mice treated with 50mg/kg Revumenib by oral gavage b.i.d. for 3 weeks. Tumor burden and immune cells were quantified by flow-cytometry.

**B.** Spleen weight.

**C.** Tumor burden in the spleen.

**D-F.** Number of myeloid (D-E) and T-cells (F) per femur (VkMYC-14551) and per femur/tibia (VkMYC-32052).

**G-H.** VkMYC-32052 cells were isolated by FACS from the bone marrow of mice after 3 weeks of revumenib treatment and subjected to RNAseq.

**G.** Volcano plot.

**H.** Barcodeplot showing RPMI8226 IRF4 knockout DEGs are enriched in revumenib treated VkMYC-32052 cells.

**I-J.** VkMYC #14551 transplanted mice were treated by oral gavage with 10 mg/kg Inobrodib q.d. with or without revumenib (50mg/kg b.i.d) for 3 weeks. Body weights during treatment period (J) and change in body weight at the end of treatment period (K).

Significance was assessed using Wilcoxon test: ****p<0.0001, ***p<0.001, **p<0.01, *p<0.05.

### References


1. Krivtsov, A.V., Evans, K., Gadrey, J.Y., Eschle, B.K., Hatton, C., Uckelmann, H.J., Ross, K.N., Perner, F., Olsen, S.N., Pritchard, T., et al. (2019). A Menin-MLL Inhibitor Induces Specific Chromatin Changes and Eradicates Disease in Models of MLL-Rearranged Leukemia. Cancer Cell *36*, 660-673.e11.
2. Uckelmann, H.J., Kim, S.M., Wong, E.M., and Armstrong, S.A. (2020). Therapeutic targeting of preleukemia cells in a mouse model of NPM1 mutant acutemyeloid leukemia. Science *367*, 586–590.
